# Magnesium isoglycyrrhizinate alleviates alcohol-associated liver disease through targeting HSD11B1

**DOI:** 10.1101/2025.10.02.679991

**Authors:** Lu Xiao, Lu Li, Shasha Wu, Zhaoyi Che, Yuyang Du, Jingyi Zheng, Jingsong Yan, Hao Wang, Hong Zhang, Yan Li, Jia Xiao

## Abstract

While magnesium isoglycyrrhizinate (MgIG) is a clinically approved therapy for alcohol-associated liver disease (ALD), its precise molecular targets and mechanisms remain uncharacterized. This study aimed to define MgIG’s hepatoprotective actions in chronic-binge ALD mouse models and ethanol/palmitic acid-exposed AML-12 hepatocytes. Through an integrated strategy encompassing RNA sequencing, molecular docking, and microscale thermophoresis, we discovered that MgIG directly binds to hydroxysteroid 11-beta dehydrogenase 1 (HSD11B1) at residue 187, a finding corroborated by molecular dynamics simulations. *In vivo*, MgIG markedly attenuated alcohol-induced liver injury, evidenced by ameliorated histological damage, reduced hepatic steatosis, and normalized liver-to-body weight ratios. *In vitro*, it effectively reduced lipid accumulation, inflammation, and apoptosis. Mechanistically, RNA sequencing identified isopentenyl diphosphate delta isomerase 1 (IDI1) as a key downstream effector. Hepatocyte-specific genetic manipulations confirmed that MgIG modulates the SREBP2-IDI1 axis, thereby suppressing lipogenesis, inflammatory responses, and apoptotic pathways. We reveal HSD11B1 as a novel direct molecular target of MgIG and elucidate its therapeutic mechanism through the HSD11B1-SREBP2-IDI1 signaling axis, which profoundly impacts ALD pathogenesis. These findings not only validate MgIG’s clinical utility but also highlight a promising new therapeutic target for ALD.

## Introduction

Alcohol-associated liver disease (ALD) encompasses a spectrum of hepatic pathologies ranging from steatosis to fibrosis, cirrhosis, and ultimately, hepatocellular carcinoma, contributing significantly to the global burden of disease and premature mortality ^1,2^. With alcohol use disorder affecting approximately 283 million individuals worldwide (5.1% of the population), alcohol emerges as a leading cause of cirrhosis, conferring a 260-fold increased risk of liver-related death ^3^. Excessive alcohol consumption, the primary etiological factor in ALD, triggers a cascade of pathological events, including oxidative stress, inflammation, and lipid metabolism dysregulation ^4^. Current ALD management primarily focuses on alcohol abstinence and supportive care. While potential therapeutic targets, such as oxidative stress pathways, inflammatory cascades, and the gut-liver axis, have been extensively investigated, the identification of druggable hepatic signaling pathways remains elusive ^5,6^.

Hydroxysteroid 11-beta dehydrogenase 1 (HSD11B1), a cortisol-regenerating enzyme predominantly expressed in the liver and kidneys, elevates endogenous glucocorticoids and has been implicated in the pathogenesis of metabolic dysfunction-associated steatotic liver disease (MASLD) and liver fibrosis ^7,8^. Similarly, isopentenyl diphosphate delta isomerase 1 (IDI1), a cytoplasmic enzyme involved in lipid metabolism, has been identified as a key risk factor for hepatocellular carcinoma ^9^ and intrahepatic cholestasis ^10^. Emerging evidence suggests that exercise may ameliorate MASLD by downregulating IDI1 expression ^11^. However, the precise roles of HSD11B1 and IDI1 in ALD progression remain to be elucidated.

Magnesium isoglycyrrhizinate (MgIG), a fourth-generation glycyrrhizin (GL) derivative and a magnesium salt of a single stereoisomer (Fig. 1A), exhibits enhanced pharmacological properties compared to its predecessors. As an 18α-GL, MgIG possesses greater lipophilicity than the β-isomer, facilitating superior binding to target cell receptors and steroid hormones ^12^. This enhanced binding capacity translates to potent anti-inflammatory effects and antioxidant effects ^13^. MgIG has demonstrated therapeutic efficacy in various liver diseases, including drug-induced liver injury (DILI) ^14^, MASLD ^15^, and liver fibrosis ^16^. Notably, MgIG has been preliminarily shown to inhibit ethanol-induced activation of the hedgehog signaling pathway in hepatocytes ^17^ and attenuate acute alcohol-induced hepatic steatosis in a zebrafish model by modulating lipid metabolism-related gene expression ^18^. Furthermore, MgIG has been recognized by the Chinese Society of Hepatology for its clinical utility in ALD treatment ^19^. Despite its therapeutic promise and proven safety, the precise molecular targets of MgIG in the liver remain poorly defined, hindering the development of optimized treatment strategies for ALD. This study employed *in vivo* and *in vitro* ALD models to investigate the hepatoprotective mechanisms of MgIG, revealing its direct interaction with HSD11B1 as a key mediator of its therapeutic effects and the involvement of downstream signaling of the sterol regulatory element binding protein 2 (SREBP2)-IDI1 axis.

**Fig. 1.**
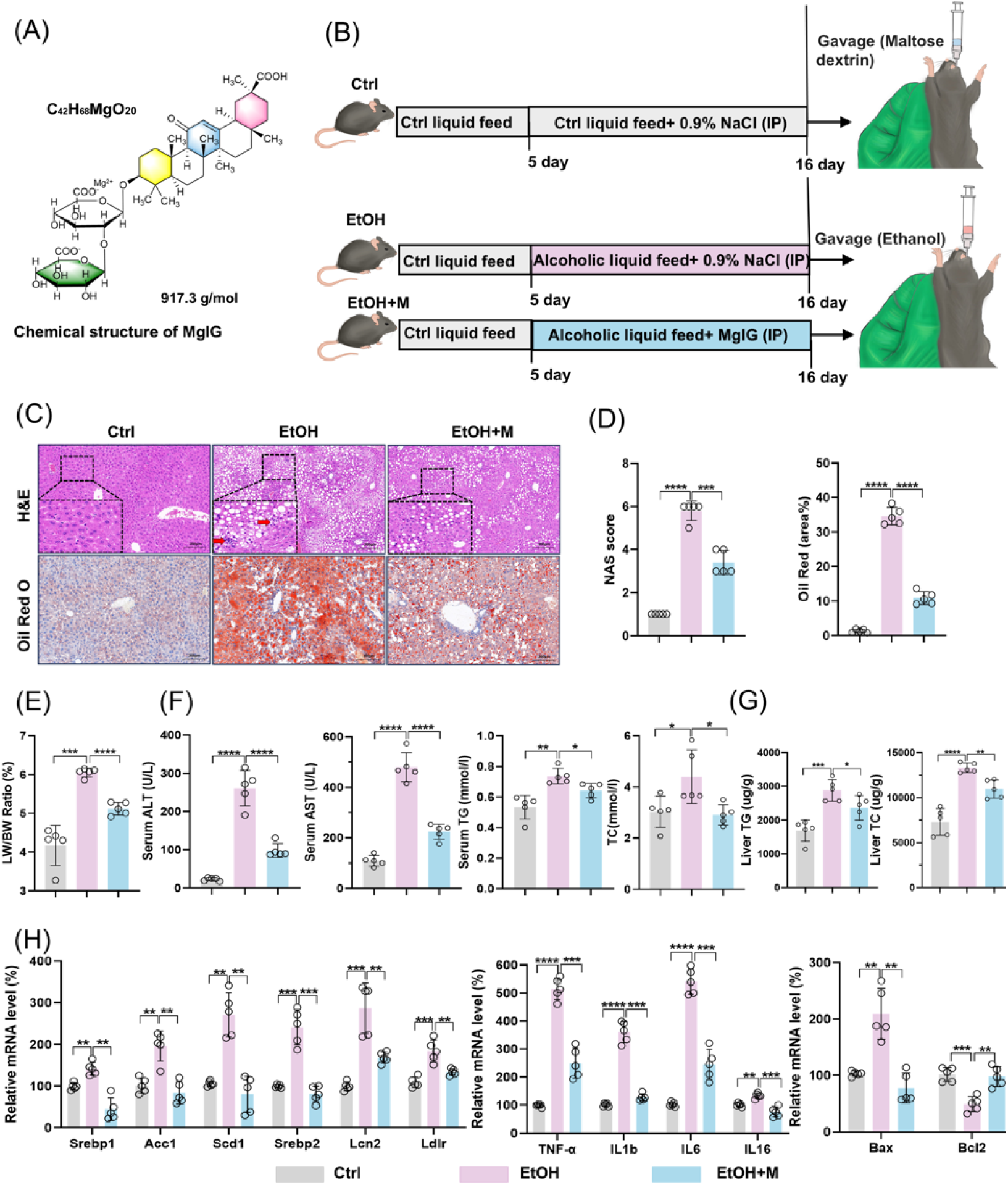
MgIG alleviates liver damage in a mouse model of chronic-binge alcoholic liver disease (the NIAAA model). (A) Chemical structure of MgIG. (B) Flowchart illustrating the modeling process for the NIAAA model. IP, intraperitoneal injections. (C) Representative results of H&E and Oil Red O staining from the livers of mice in Ctrl, EtOH and EtOH+M groups (n = 5). (D) Alterations in NAS (NAFLD activity score) and Oil Red O quantification (a.v.: arbitrary value) (n = 5). (E) Ratios of liver weight to body weight (LW/BW) in mice (n = 5). (F) Alterations in serum biochemical parameters (ALT, AST, TG, and TC) in mice in three groups (n = 5). (G) Alterations in liver parameters (TG and TC) in mice in three groups (n = 5). (H) Alterations in mRNA expression of lipid metabolism genes (*Srebp1*, *Srebp2*, *Acc1*,*Scd1, Lcn2, and Ldlr*), systemic inflammation markers (*Tnf-*_α_, *Il1b, Il-6,* and *Il-16*), and apoptosis-related genes (*Bax* and *Bcl2*) in the mice liver. The data are presented as mean ± SD. \**P* < 0.05, \*\**P* < 0.01, \*\*\**P* < 0.001, \*\*\*\**P* < 0.0001. Scale bar: 50 μm.

## Methods

### Chemicals and reagents

MgIG was generously provided by Chia-Tai Tianqing Pharmaceutical Group Co., Ltd. (Nanjing, China) in the form of a magnesium isoglycyrrhizinate injection (5 mg/mL). High-purity ethanol was obtained from MACKLIN (Shanghai, China). Palmitic acid (PA) and fatty acid-free bovine serum albumin (BSA) were purchased from Sigma-Aldrich (St. Louis, MO, USA). All cell culture reagents and consumables were obtained from Gibco (Carlsbad, CA, USA) or Corning Incorporated (Corning, NY, USA). Primary antibodies against tumor necrosis factor-α (TNF-α, #11948), interleukin-6 (IL-6, #12912), glyceraldehyde-3-phosphate dehydrogenase (GAPDH, #2118S), Bax (#2772S), Bcl-2 (#3498S), and hemagglutinin (HA, #3724S) were purchased from Cell Signaling Technology (Shanghai, China). Antibodies against IDI1 (#11166-2-AP) and SREBP2 (#28212-1-AP) were obtained from Proteintech (Wuhan, China). An antibody against HSD11B1 (#AF3397) was purchased from R&D Systems (Minneapolis, MN, USA). A Flag antibody (F1804) was purchased from Sigma-Aldrich. All antibody validation experiments were performed according to the manufacturers’ protocols. Protein A/G Magarose beads (#80105G) were purchased from Invitrogen (Carlsbad, CA, USA).

### Animal experiments

Male C57BL/6 wild-type mice (7 weeks old; 19-21 g) were purchased from Guangzhou Qingle Biosciences Co., Ltd. (Guangzhou, China). Mice were acclimated to the housing environment for one week prior to experimental procedures. Alcoholic liver injury was induced using a modified National Institute on Alcohol Abuse and Alcoholism (NIAAA) model ^20^. Randomization was performed by an investigator not involved in subsequent data collection and analysis using a random number table. Briefly, mice (n = 5 per group) were initially fed *ad libitum* with the control (Ctrl) Lieber-DeCarli liquid diet for 5 days to facilitate acclimation. Subsequently, the ALD groups (designated as ’EtOH’ and ’EtOH+M’ in figures) received the Lieber-DeCarli diet containing 5% (v/v) ethanol for 10 days, while the control group (designated as ’Ctrl’ in figures) received an isocaloric control diet. Body weight and food intake were monitored daily. On day 11, ethanol-fed mice and their pair-fed controls were administered a single dose of ethanol (5 g/kg body weight) or an isocaloric maltose dextrin solution, respectively, via oral gavage in the early morning. Mice were euthanized 9 h later (Fig. 1B). A vehicle-MgIG control group was not included in this study because: (1) MgIG is a clinically approved drug with an established safety profile; (2) the dosage used in this study was calculated based on human clinical dosages using a standard conversion formula; and (3) the vehicle safety of MgIG with similar dose has been previously confirmed in an animal study ^21^. For viral infections, mice (n = 5 per group) received tail vein injections of 5 × 10^11^ genome copies of either AAV8 control or AAV-shRNA/plasmid. After 14 days, mice were fasted for 4 h at the end of the dark cycle and then euthanized to confirm hepatic upregulation or downregulation of *Hsd11b1*, *Srebp2*, or *Idi1*. These mice were then subjected to the NIAAA model and/or MgIG administration as described above. Target gene expression in the liver was verified using quantitative PCR and Western blotting. All experimental procedures were approved by the Ethical Committee of Jinan University, China (IACUC-20220708-04).

### RNA sequencing and functional enrichment analysis

Total RNA was extracted from liver tissue samples using TRIzol reagent (Invitrogen) following the manufacturer’s instructions. RNA quality was assessed using a Nanodrop ONE spectrophotometer (Thermo Fisher, Waltham, MA, USA). Samples with A260/A280 ratios above 1.8 and A260/A230 ratios above 2.0 were deemed acceptable for further processing. RNA integrity was evaluated using an Agilent 2200 TapeStation (Agilent Technologies, Santa Clara, CA, USA), and only samples with an RNA integrity number above 7.0 were used for library preparation. Ribosomal RNA (rRNA) was depleted from total RNA using the Epicentre Ribo-Zero rRNA Removal Kit (Illumina) and fragmented to an average size of 200 bp. First-strand and second-strand cDNA synthesis, followed by adapter ligation and low-cycle enrichment, were performed using the NEBNext Ultra RNA Library Prep Kit for Illumina (New England Biolabs, Ipswich, MA, USA). Library quality was assessed using the Agilent 2200 TapeStation and Qubit 2.0 fluorometer (Thermo Fisher). Libraries were diluted to 10 pmol/L and subjected to cluster generation on a pair-end flow cell, followed by sequencing (2 × 150 bp) on an Illumina HiSeq 3000. Raw sequencing reads were processed to remove adapter sequences, low-quality reads, and reads containing poly-N stretches. Clean reads were aligned to the mouse reference genome using HISAT2 with default parameters. Gene expression levels were quantified using HTSeq, and differential expression analysis was performed using DESeq with read counts as input. The Benjamini-Hochberg method was used to adjust p-values for multiple testing. Differentially expressed genes (DEGs) were defined as those with a fold change > 2 and an adjusted p-value < 0.05. Heatmap visualization and Kyoto Encyclopedia of Genes and Genomes (KEGG) pathway enrichment analysis were performed using the identified DEGs. A p-value < 0.05 was considered statistically significant for KEGG pathway enrichment.

### Biochemical and cytokine analyses of mice serum and cell supernatants

Serum or hepatic levels of alanine aminotransferase (ALT), aspartate aminotransferase (AST), triglyceride (TG) and total cholesterol (TC) were measured using BK2800 (BIOBASE, Jinnan, China). Cell supernatants cytokine levels were determined by using corresponding ELISA kits from R&D Systems (TNF-_α_: #VAL609; IL-6: #VAL604G).

### Liver tissue histology

Liver tissue samples were fixed in 10% neutral buffered formalin and subsequently embedded in paraffin. Tissue sections (5 μm thickness) were prepared and stained with hematoxylin and eosin (H&E) and Oil Red O for histological examination. Images were acquired using a LEICA Qwin Image Analyzer (Leica Microsystems, Milton Keynes, UK). The nonalcoholic fatty liver disease activity score (NAS) was determined for each group as previously described ^22^.

### Cell culture and transfection

The AML-12 mouse normal hepatocyte cell line was obtained from the Cell Bank of Type Culture Collection, Chinese Academy of Sciences (Shanghai, China). Cells were cultured in Dulbecco’s modified Eagle’s medium supplemented with 10% (v/v) fetal bovine serum, incubated at 37 °C with 5% CO_2_ in a cell incubator. To induce an ALD-like phenotype in AML-12 cells, cells were exposed to a combination of 25 mmol/L ethanol and 0.1-1 mmol/L palmitic acid (PA; Sigma-Aldrich; #P0500) for 24 h to determine the optimal concentration for inducing cellular injury, as previously described (12). We chose the combination of ethanol and palmitic acid in the *in vitro* experiments because ethanol-only treatment could not phenocopy alcoholic cell death, inflammation, oxidative stress, and the dysregulated lipid metabolism in hepatocytes exhibited in early-stage ALD ^23–26^. Following this initial treatment, varying concentrations of MgIG were added to the culture medium for an additional 24-hour to assess its effects and determine the optimal dose. For gene knockdown and overexpression experiments, Lipofectamine 3000 reagent (Thermo Fisher; #L3000001) was used to transfect AML-12 cells with *Hsd11b1* or *Idi1*-targeting constructs in 6-well plates.

### Cell lipid accumulation, viability, and apoptosis assays

Intracellular lipid droplets were visualized by staining AML-12 cells with Nile Red. Briefly, cells were washed with phosphate-buffered saline (PBS), fixed with 4% formaldehyde for 10 min, and then stained with Nile Red solution (0.1 mg/mL; #N8440, Solarbio, Beijing, China) for 10 min at room temperature in the dark. Cells were then counterstained with DAPI (5 μg/mL; Sigma-Aldrich; D9542) for 5 min. Cell viability and apoptosis were assessed in 96-well plates using a CCK8 cell viability assay and a lactate dehydrogenase (LDH) cytotoxicity assay kit, respectively, according to the manufacturer’s instructions (MedChem Excess, Shanghai, China; HY-K1090 and HY-K0301).

### Quantitative PCR

Total RNA was extracted from liver tissue or cultured cells using TRIzol reagent. The first strand cDNA was synthesized from total RNA using a PrimeScript RT Reagent Kit (Takara, Shiga, Japan; #RR047A) following the manufacturer’s instructions. Quantitative PCR reaction was performed with the SYBR Premix Taq Quantitative PCR Kit (Takara) according to the manufacturer’s instructions, on a qTOWER3 machine (Analytik Jena AG, Jena, Germany). Primer information is as listed in Supporting Information Table S1. β-actin was used as an internal control. The relative quantification of mRNA expression levels was calculated using the 2^−ΔΔCt^ method. All qPCR experiments were conducted in compliance with the MIQE guidelines ^27^.

### Protein extraction and Western blot

Samples were lysed using Pierce RIPA buffer (Thermo Fisher) and their protein concentrations were measured with the Bio-Rad Protein Assay (Bio-Rad, Hercules, CA). Western blot analyses of all proteins were conducted with β-actin serving as the internal control.

### Coimmunoprecipitation (Co-IP)

Co-IP buffer (50 mM Tris-HCl, 5 mM EDTA, 150 mM NaCl, and 1% NP-40 pH 7.6) mixed with protease inhibitor cocktail (Roche, Bern, Switzerland) were used to lyse the cells. The samples were then incubated with the corresponding antibodies and Protein A/G Magarose beads at 4°C overnight.

### Protein structure modeling and molecular docking

We first employed DecoupleR’s ^28^ built-in Univariate Linear Model (ULM) to compute the transcription factor activity perturbed by MgIG, based on the expression of differentially expressed genes. The findings revealed that *Srebf2* was the transcription factor most significantly reduced by MgIG. We then identified potential SREBP2/IDI1-interacting proteins using the STRING and PubMed databases. The 3D structures of these proteins were obtained from the Protein Data Bank (PDB) or UniProt, while the 3D structure of MgIG was retrieved from PubChem. Molecular docking simulations were performed using Schrödinger software to predict the binding affinity (Glide score) of MgIG to each protein. The lowest-energy docking conformations were visualized using PyMOL.

### Microscale thermophoresis (MST) assay

The binding interaction between MgIG and HSD11B1 was assessed using a Monolith NT.115 Blue/Green instrument (NanoTemper Technologies, Munich, Germany). Initially, we evaluated the affinity and labeling efficiency of the dye with the His-tagged protein. Subsequently, HSD11B1 proteins (100 nM) were labeled and incubated with MgIG at a concentration gradient ranging from 50 µM to 1.53 nM for 30 min at room temperature. The samples were then analyzed at 60% MST power and a temperature of 25°C. Dissociation constants (Kd) were calculated based on a 1:1 binding stoichiometry.

### Molecular dynamics (MD) simulations

Three molecular systems were constructed in this study: first, Compound-HSD11B complex was established through molecular docking. Second, the last frame from 100 ns MD simulation was used to generate APO-HSD11B1 by removing the ligand. Finally, based on this conformation and reported portal vein ethanol concentrations in ALD, five ethanol molecules (0.1 g/dL) were incorporated to construct the EtOH-HSD11B1 system, simulating the ALD environment ^29–31^. For each system, receptor was prepared and missing atoms of residues were fixed by the advanced PDB-Preparation tool in Yinfo Cloud Computing Platform using PDBFixer and the tLEaP module in AmberTools 20. The AM1-BCC charges were calculated for compound magnesium isoglycyrrhizinate by the Amber antechamber program ^32^. MD simulations was performed using AmberTools 20 package with AMBER ff19SB ^33^ and GAFF ^34^ force field.

The system was solvated by a truncated octahedron water box using OPC water model with a margin of 10 Å. Periodic boundary condition (PBC) was used and the net charge neutralized by Na^+^ ions. Nonbonded van der Waals interactions were calculated using the Lennard-Jones 12-6 potentials with a 10 Å cutoff, while long-range electrostatics were treated using the Particle Mesh Ewald (PME) algorithm. The SHAKE algorithm was applied to constrain bonds involving hydrogen atoms ^35^. To removed improper atom contacts, the structure was first minimized by (1) 2500 steps of steepest descent and 2500 steps of conjugate gradient, under a harmonic constraint of 10.0 kcal/(mol·Å2) on heavy atoms; (2) relaxing the entire system by 10000 steps of steepest descent and 10000 steps of conjugate gradient. And then the system was gradually heated up to 300 K by a 20 ps NVT simulation. Subsequently, two steps of equilibration phases were carried out: (1) a 200 ps NPT simulation with constraints on heavy atoms followed by (2) a 1 ns NVT simulation without restraint. The temperature was maintained at 300 K using the Berendsen thermostat with 1 ps coupling constant and the pressure at 1 atm using Monte Carlo barostat with 1 ps relaxation time. Finally, the system was subjected to a 100 ns NVT simulation with a time step of 2 fs. The root-mean-square deviation (RMSD), root-mean-square fluctuation (RMSF), radius of gyration (RG), solvent-accessible surface area (SASA) and hydrogen bonds were analyzed by the CPPTRAJ module, while principal component analysis (PCA) and dynamic cross-correlation matrix (DCCM) were analyzed by R package Bio3D ^36^. The binding free energies were calculated using the Molecular Mechanics Generalized Born (MM/PBSA) method for the 100 ns MD trajectory ^37^.

### HSD11B1 enzyme activity assays

The enzyme activity of HSD11B1 in cells was evaluated according to its capability to convert cortisone to cortisol ^38^. AML-12 cells were seeded into 6-well culture plates and induced ALD as above. Different concentrations of MgIG (0.1, 0.25, and 0.5 mg/mL) were added 24 h prior to the co-treatment, with a control group included. AML-12 cells were exposed to 160 nmol/L cortisone for 24 h. The reaction mixtures were collected and analyzed for cortisol levels using an ELISA kit (R&D Systems, Minneapolis, MN, USA), according to the manufacturer’s instructions.

### Statistical analysis

The data from each group are presented as the mean ± standard deviation (SD). For data that followed a normal distribution, statistical comparisons between two groups were conducted using an unpaired two-tailed Student’s t-test; for comparisons involving three or more groups, a two-way ANOVA followed by a Student-Newman-Keuls post hoc test was applied (Prism 5.0, Graphpad Software, Inc., San Diego, CA). A P-value of less than 0.05 was deemed statistically significant.

## Results

### MgIG alleviates liver injury in a mouse model of ALD

To investigate the hepatoprotective effects of MgIG *in vivo*, we employed a well-established chronic-binge NIAAA mouse model of ALD (Fig. 1B). Based on the drug’s clinical instructions, previous studies ^21,39^, and our preliminary experimental results (including body weight change, serum ALT/AST, hepatic TG and TC levels, and histological NAS scores) (Fig. S1), we determined an optimal MgIG dose of 50 mg/kg. Compared to mice fed a normal diet (Ctrl group), mice in the alcohol-fed control group (EtOH) exhibited characteristic histological features of ALD, including steatosis and inflammatory cell infiltration, resulting in an elevated NAS. MgIG treatment (EtOH+M) effectively attenuated these pathological changes (Fig. 1C and 1D). Consistent with these histological findings, the liver-to-body weight ratio, a marker of liver injury, was significantly increased in the EtOH group, and this increase was markedly reduced by MgIG treatment (Fig. 1E). Notably, significant differences were observed between the EtOH group and the MgIG-treated (EtOH+M) group in serum levels of liver enzymes (ALT and AST), serum lipid parameters (TG and TC), as well as Liver TG and TC contents-key indicators of liver function and lipid metabolism. (Fig. 1F). In line with the observed histological and physiological improvements, MgIG treatment also reduced the expression of genes involved in lipid synthesis metabolism (*Srebp1*, *Srebp2*, *Acc1*, *Scd1, Lcn2, and Ldlr*), inflammation (*Tnf-*_α_ and *Il-6*), and pro-apoptosis (*Bax*) while restored the level of anti-apoptotic gene (*Bcl2*) in the liver tissue of EtOH mice (Fig. 1G-1H).

### MgIG protects against ethanol-induced hepatocyte injury via IDI1

To further elucidate the molecular mechanisms underlying the hepatoprotective effects of MgIG, we performed RNA sequencing analysis on liver tissue from the NIAAA mouse model. DEG analysis revealed a significant upregulation of *Idi1* in the EtOH group compared to the Ctrl group (Fig. 2A, Fig. S4A-4B). Furthermore, KEGG pathway enrichment analysis identified metabolic pathways among the top five downregulated pathways in the MgIG-treated group compared to the EtOH group, with *Idi1* consistently ranked among the top two genes in these pathways (Fig. 2C). To further investigate the role of IDI1 and its potential interaction with MgIG, we established an *in vitro* model of ethanol/palmitic acid (EtOH/PA)-induced hepatocyte injury using AML-12 cells. We first optimized the EtOH/PA concentrations to induce significant cell death and lipid accumulation (Fig. S2), identifying 250 mmol/L ethanol and 0.2 mmol/L PA as the optimal combination (Fig. S2A-2C). MgIG treatment (0.1-1.0 mg/mL) demonstrated a dose-dependent protective effect against EtOH/PA-induced injury, as evidenced by improved cell viability, reduced lipid accumulation, and decreased apoptosis (Fig. 2D-2G, Fig. S2D). Consistent with the *in vivo* findings, EtOH/PA treatment increased the expression of genes involved in lipid metabolism (*Srebp1*, *Acc1*, *Scd1, Srebp2, Lcn2,* and *Ldlr*) and inflammation (*Tnf-*_α_ and *Il-6*), and MgIG treatment effectively reversed these effects (Fig. 2H). Based on these results, we selected 0.25 mg/mL as the optimal MgIG concentration for subsequent *in vitro* experiments.

**Fig. 2.**
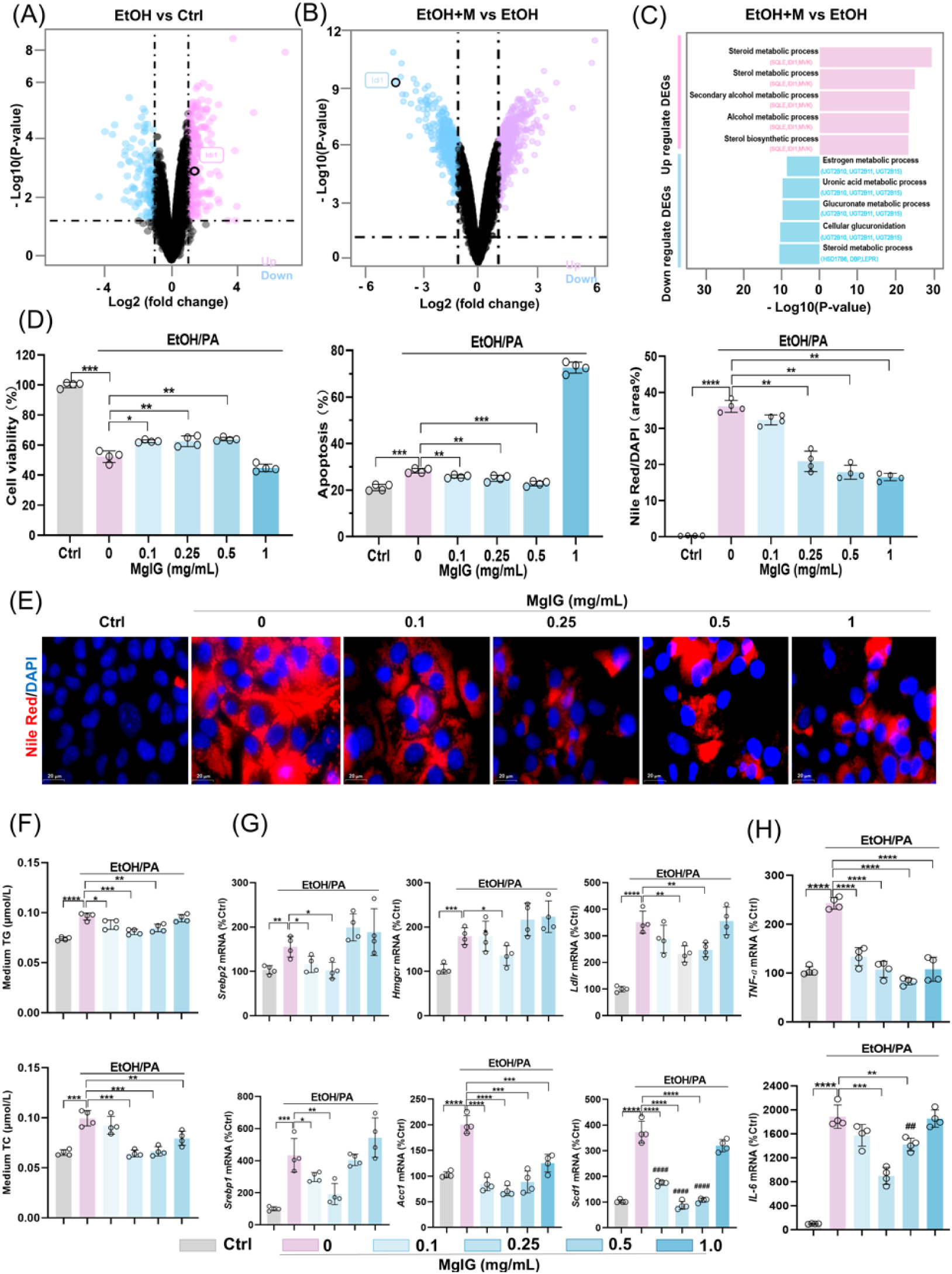
MgIG protects ethanol-induced hepatocyte injury in a cell model. (A) Volcano plot analysis showing differentially expressed genes between the EOH group and the Ctrl group after RNA-seq in the liver of mice. (B) Volcano plot analysis showing differentially expressed genes between the EtOH+M and the EtOH group after RNA-seq in liver of mice. (C) The top 5 regulated pathways (both up- and down-regulated) and the top 3 genes within each pathway are shown (EtOH+M vs EtOH).. (D) Changes in AML-12 cell viability, apoptosis, and Nile Red staining signals after ethanol/PA treatment, with or without co-treatment with different MgIG doses (EtOH+M) (0, 0.1, 0.25, 0.5, and 1 mg/mL) (n = 4). (E) Representative images of Nile Red staining in AML-12 cells treated with ethanol/PA, with or without co-treatment with different MgIG doses. (F)Quantification of triglyceride (TG) and total cholesterol (TC) levels in the in the culture supernatant of AML-12 cells treated with EtOH/PA in the presence or absence of increasing concentrations of MgIG (n = 4). (G-H) Changes in mRNA expression of lipid metabolism genes (*Srebp1, Acc1*,*Scd1, Srebp2, Lcn2,* and *Ldlr*) and systemic inflammation markers (*Tnf-*_α_ and *Il-6*) in AML-12 cells treated with ethanol/PA, with or without co-treatment with different MgIG doses. Data are expressed as mean ± SD. \**P* < 0.05, \*\**P* < 0.01, \*\*\**P* < 0.001. Scale bar: 20 μm.

In line with the *in vivo* observations, *Idi1* expression was significantly increased in AML-12 cells following EtOH/PA treatment, and this effect was attenuated by MgIG treatment (Fig. 3A and S3A-3C). As shown in Fig. 3B and 3C, transfection with siRNA and plasmid effectively decreased or increased *Idi1* mRNA levels without affecting cell viability, respectively. Knockdown of *Idi1* significantly ameliorated EtOH/PA-induced cell injury, as evidenced by reduced apoptosis, inflammatory cytokine production, and lipid accumulation. In contrast, Idi1 overexpression tended to exacerbate these pathological changes (Fig. 3D-3H, Fig. S3). Although the effects of *Idi1* overexpression on apoptosis and inflammatory cytokines did not reach statistical significance in Western blot analyses, Idi1 overexpression markedly attenuated the hepatoprotective effects of MgIG (Fig. 3E, Fig. S3A) . One possible explanation is that *Idi1* functions as a permissive factor in EtOH/PA-induced hepatocellular injury, rather than acting as a sole or rate-limiting determinant. In this context, modulation of *Idi1* may be necessary to facilitate lipid metabolic reprogramming and stress responses, but is not sufficient on its own to fully dictate cell fate decisions such as apoptosis or survival. This interpretation is consistent with the multifactorial nature of ALD, which involves the coordinated interplay of oxidative stress, inflammatory signaling, and metabolic dysregulation ^5,6^. Given that MgIG has been reported to exert broad anti-inflammatory and antioxidant effects^13^, it is likely that its hepatoprotective actions are mediated through multiple parallel pathways, with *Idi1* representing one important, but not exclusive, target. Taken together, these findings suggest that MgIG protects against ethanol-induced hepatocyte injury, at least in part, through the modulation of IDI1.

**Fig. 3.**
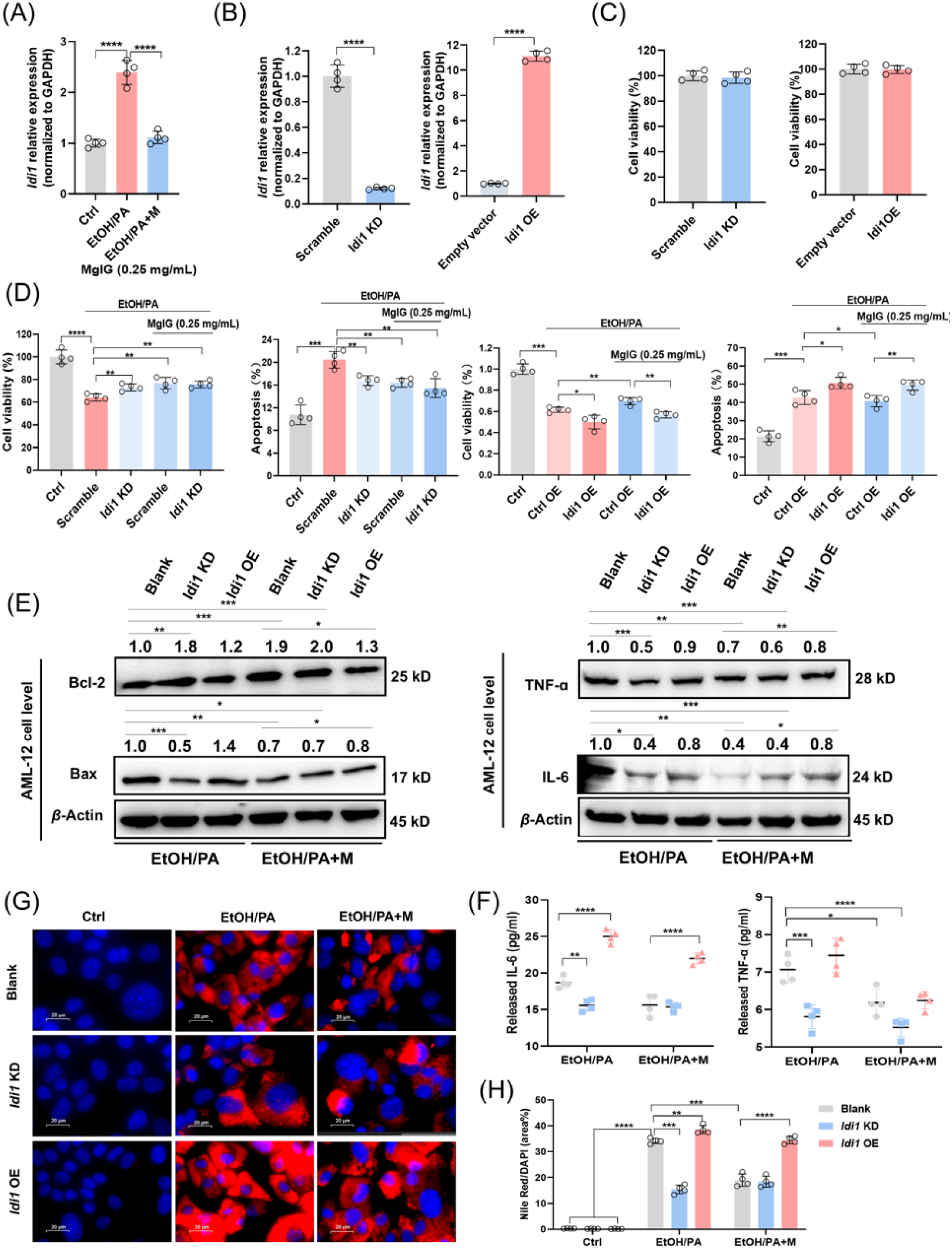
IDI1 is involved in MgIG-mediated hepatocyte protection against ethanol. (A) Quantitative PCR validation of RNA-seq results revealed changes in *Idi1* gene expression in AML-12 cells treated with ethanol/PA, with or without 0.25 mg/mL MgIG co-treatment (EtOH/PA and EtOH/PA+M) (n = 4). (B) Quantitative PCR results confirmed the knockdown or overexpression efficiency of *Idi1* genes by siRNA or *Idi1* plasmid in AML-12 cells. (C) The changes in AML-12 cell viability following *Idi1* siRNA or *Idi1* plasmid transfection (24 h) were also assessed (n = 4). (D) Changes in cell viability and apoptosis ratios were assessed in ethanol/PA-treated AML-12 cells, with or without 0.25 mg/mL MgIG, following *Idi1* knockdown/overexpression (n = 4). (E, F) Western blot results for TNF-α, IL-6, Bax, and Bcl-2 and cell supernatant results for TNF-α and IL-6 in AML-12 cells treated with ethanol/PA (EtOH/PA) and ethanol/PA + MgIG (EtOH/PA+M), with or without *Idi1* knockdown/overexpression. Numbers above the lanes indicate the mean relative density normalized to the loading control for each group (n = 3). (G, H) Nile Red staining area (%) with corresponding representative cell staining images. Data are expressed as mean ± SD. For Western blot quantification: \**P* < 0.05, \*\**P* < 0.01, \*\*\**P* < 0.001. Scale bar: 20 μm.

### MgIG directly binds to HSD11B1

To identify potential direct binding partners of MgIG, we initially performed molecular docking simulations between MgIG and IDI1. However, the resulting Glide score of -2.74 indicated a lack of favorable binding interactions, which implied that there should be an immediate binding protein in the upstream of IDI1. To identify potential upstream regulators of *Idi1* expression, we utilized the ULM within the DecoupleR package to analyze transcription factor activity. This analysis identified SREBP2 as the most significantly affected transcription factor (Fig. 4A). We then expanded our search to include upstream regulators of IDI1 identified using the STRING database. Among these proteins, HSD11B1 exhibited the highest binding affinity for MgIG, with a Glide score of -8.75 (Fig. 4B). Given the homology between human and mouse *Hsd11b1* and the need for future translational research, we focused on validating the interaction between human HSD11B1 and MgIG. We conducted 100 ns MD simulations for Compound-HSD11B1 (MgIG-HSD11B1 binding in physiological condition) and EtOH-HSD11B1 (MgIG-HSD11B1 binding in alcoholic condition), with free HSD11B1 as a control. The results revealed that HSD11B1 exhibited high flexibility and dynamic motion in its unbound state but became significantly more rigid and stable upon binding to MgIG, particularly at residues 219–233 (Fig. 4C-4E). This structural stabilization was further supported by the radius of gyration curves, which indicated a more compact conformation upon binding (Fig. S5 and 5B). Further analyses, including PCA and DCCM, demonstrated alterations in motion direction, dynamic patterns, and residue correlations. Specifically, upon binding, HSD11B1 exhibited reduced motion amplitude and weaker residue correlations, with its dynamic patterns becoming more disordered (Fig. S5C-5E). In the Compound-HSD11B1 system (Fig. 4E), key interactions included hydrogen bonds (Tyr177, Tyr183), a salt bridge (Lys187) and hydrophobic interactions (Leu126, Leu217, Ala223, Ala226, and Ile230). In contrast, the EtOH-HSD11B1 system (Fig. 4F) exhibited hydrogen bonds (Thr124, Ser170, Tyr183, Leu217, and Arg269) along with an additional salt bridge (Tyr183 and Arg269). Notably, the EtOH-HSD11B1 system demonstrated stronger hydrogen bonding and more favorable binding free energy compared to the Compound-HSD11B1 system (Fig. 4G and S5F), suggesting enhanced stability and binding affinity in an ethanol environment. Based on molecular docking and MD results, we further validated the binding between MgIG and the human full-length wild-type (WT) HSD11B1 protein as well as its site-directed mutants (Tyr177, M1; Tyr183, M2; Lys187, M3) using MST. The Kd value of the WT protein was 6.35 μM (Fig. 4H), while those of M1, M2, and M3 were 48.6 μM, 11.5 μM, and 135.6 μM, respectively (Fig. 4I). These data indicate that MgIG primarily exerts its anti-ALD effect by inhibiting HSD11B1 through the critical Lys187 site.

**Fig. 4.**
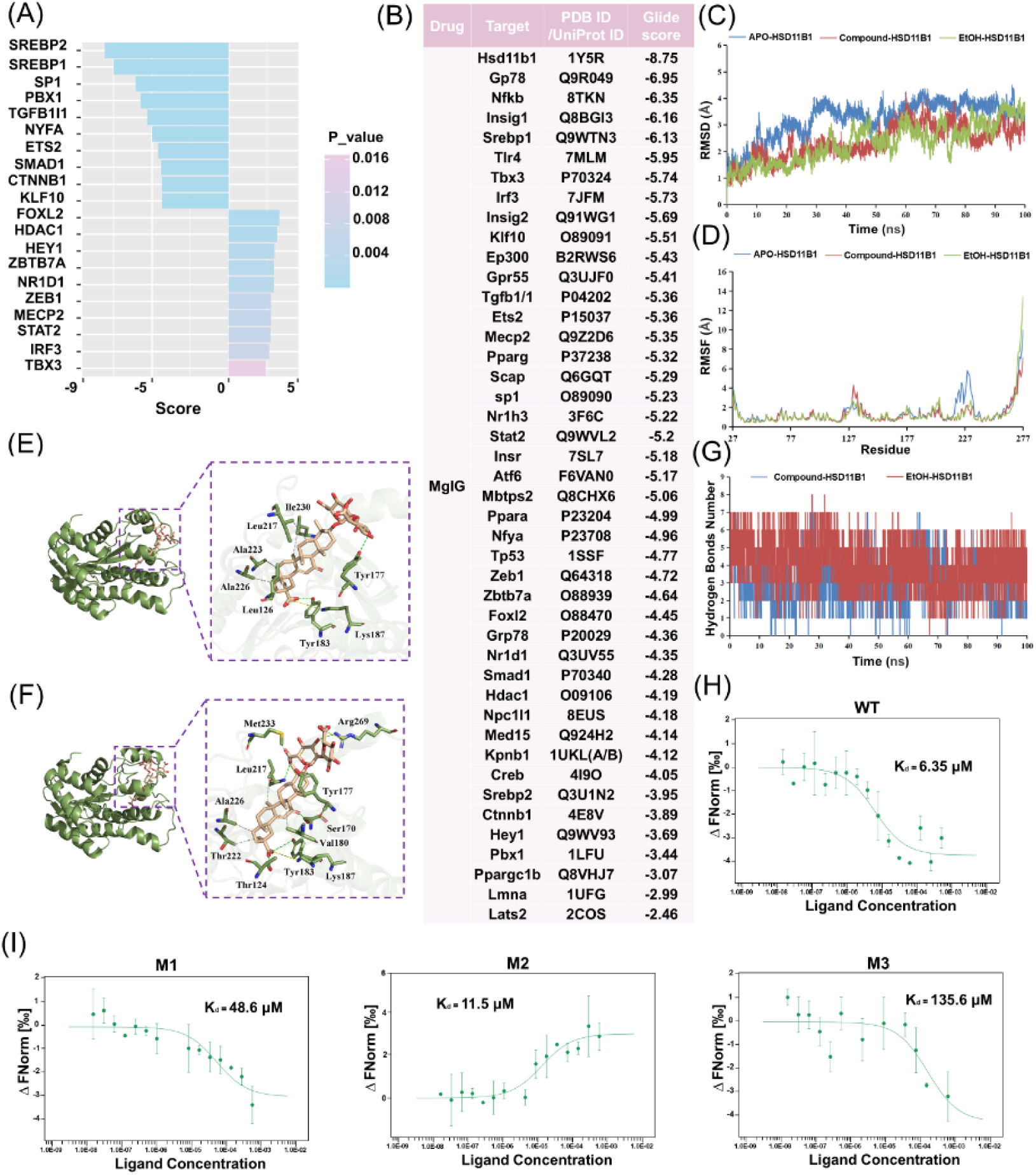
HSD11B1 is the direct binding protein of MgIG for the protection of ethanol-induced hepatocytes injury. (A) Bar chart displaying the top 10 upregulated and downregulated transcription factor activity scores in the EtOH+M group compared to the EtOH group. (B) The glide score of MgIG binding to the protein structure as determined by molecular docking analysis. (C, D) RMSD and RMSF analysis of the three systems: APO-HSD11B1 (blue) represents unbound HSD11B1 in a physiological saline system, Compound-HSD11B1 (red) represents HSD11B1 bound to MgIG in a physiological saline system, and EtOH-HSD11B1 (green) represents HSD11B1 bound to MgIG in a 0.1 mg/mL ethanol solvent system. (E, F) Molecular modeling analysis of MgIG binding at the HSD11B1 domain in normal saline and ethanol systems. Left: Cartoon view of MgIG at the HSD11B1 binding site. Right: Close-up surface view of MgIG at the HSD11B1 binding sites. (G) Hydrogen bond analysis of HSD11B1-MgIG interactions in normal saline and ethanol systems. (H, I) The microscale thermophoresis (MST) assay demonstrated direct binding between varying doses of MgIG and human HSD11B1 protein at residues 187. WT: wild-type HSD11B1; M1, M2, M3: point mutations at Tyr177, Tyr183, and Lys187, respectively.

To further investigate the functional significance of this interaction, we modulated *Hsd11b1* expression in AML-12 cells using siRNA- and plasmid-mediated knockdown and overexpression, respectively. As shown in Fig. 5A-5C, transfection with siRNA and plasmid effectively decreased or increased *Hsd11b1* mRNA levels without affecting cell viability, respectively. Knockdown of *Hsd11b1* significantly attenuated EtOH/PA-induced inflammation, apoptosis, and lipid accumulation, even in the absence of MgIG treatment. Conversely, overexpression of *Hsd11b1* exacerbated these effects. Importantly, MgIG treatment partially reversed the detrimental effects of *Hsd11b1* overexpression (Fig. 5D-5F, Fig. S4C, 4F, 4G). To further investigate whether the binding of MgIG to HSD11B1 affects its enzymatic activity, we added different concentrations of MgIG (0.1, 0.25, and 0.5 mg/mL) to the cellular model. ELISA results showed that HSD11B1 activity was significantly elevated in the ALD group, whereas increasing concentrations of MgIG led to a dose-dependent reduction in HSD11B1 activity (Fig. 5G).Given the low basal activity under normal conditions, we further examined cortisol levels in the supernatant of AML-12 cells overexpressing *Hsd11b1*, with or without MgIG co-treatment (Fig. 5H). These findings suggested that MgIG exerted its protective effects against EtOH/PA-induced hepatocyte injury, at least in part, by directly binding to and modulating the activity of HSD11B1.

**Fig. 5.**
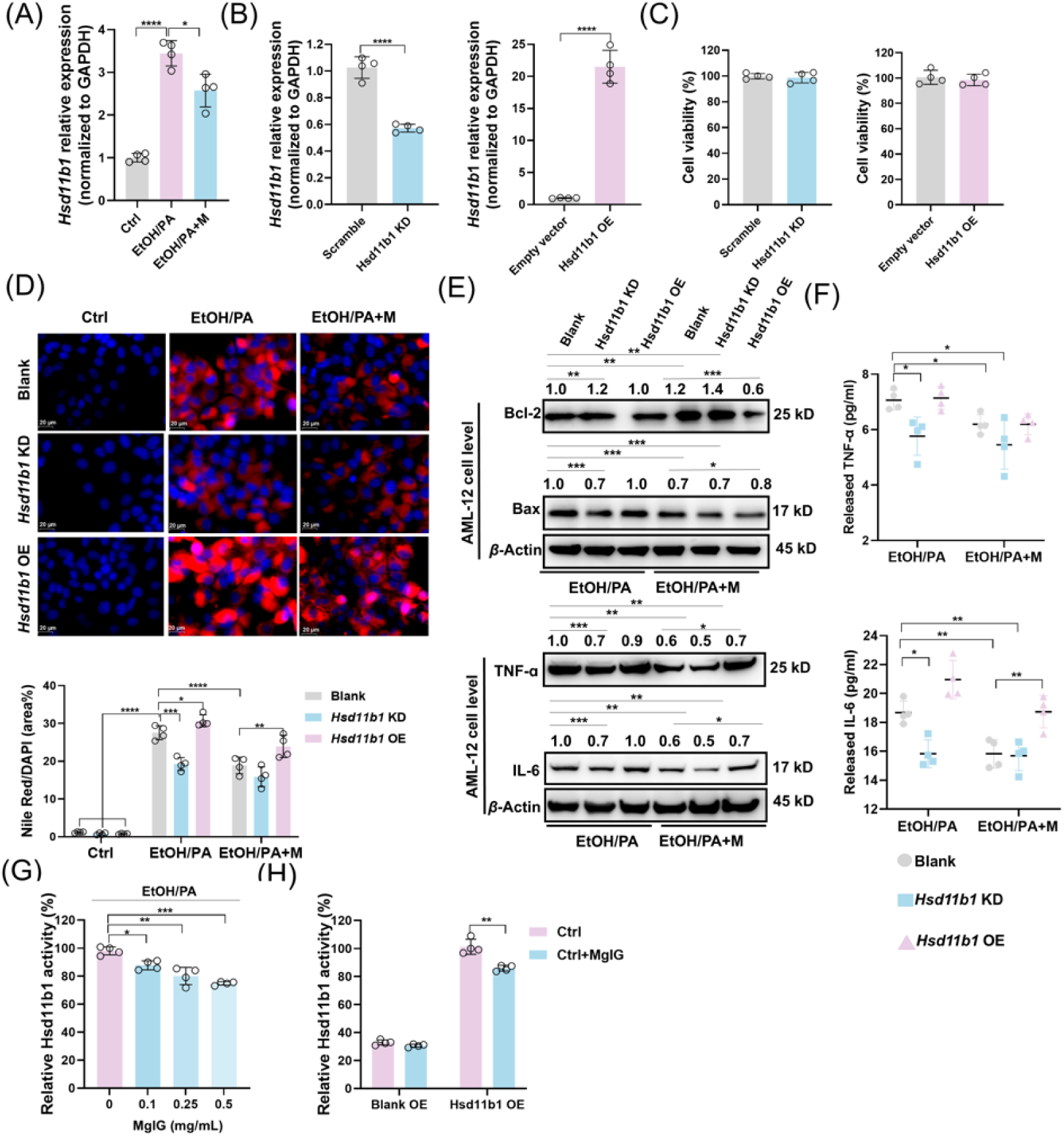
HSD11B1 is involved in MgIG-mediated hepatocyte protection against ethanol. (A) Quantitative PCR validation of RNA-seq results revealed changes in *Hsd11b1* gene expression in AML-12 cells treated with ethanol/PA, with or without 0.25 mg/mL MgIG co-treatment (EtOH/PA and EtOH/PA+M) (n = 4). (B, C) Quantitative PCR confirmed the knockdown efficiency of *Hsd11b1* (siRNA) and the overexpression efficiency of *Hsd11b1*(plasmid) in AML-12 cells, and cell viability changes were assessed 48 h after transfection (n = 4). (D) Nile Red staining area (%) with corresponding representative cell staining images (n=4). (E, F) Western blot results for TNF-α, IL-6, Bax, and Bcl-2 and cell supernatant results for TNF-α and IL-6 in AML-12 cells treated with ethanol/PA (EtOH/PA) and ethanol/PA + MgIG (EtOH/PA+M), with or without *Hsd11b1* knockdown/overexpression. Numbers above the lanes indicate the mean relative density normalized to the loading control for each group (n = 3). (G,H) Impact of MgIG on Hsd11b1 Activity. Data are expressed as mean ± SD. \**P* < 0.05, \*\**P* < 0.01, \*\*\**P* < 0.001. Scale bar: 20 μm.

### MgIG protects hepatocyte partially through the HSD11B1–SREBP2–IDI1 axis

To investigate the relationship between SREBP2 and HSD11B1, we conducted co-IP experiments, which revealed a direct interaction between the precursor form of SREBP2 (p-SREBP2) and HSD11B1 (Fig. S4E-4G). This interaction appeared to be reduced following MgIG treatment. (Fig. 6A and 6B). To further explore this interaction, we modulated *Hsd11b1* expression in EtOH/PA induced AML-12 cells with or without MgIG treatment (Fig. 6C-6D, Fig. S6). Knockdown of *Hsd11b1* reduced nuclear translocation of SREBP2 (n-SREBP2), while *Hsd11b1* overexpression promoted the accumulation of n-SREBP2, an effect further enhanced by the EtOH/PA treatment. These findings suggest that HSD11B1 may regulate the processing and nuclear translocation of SREBP2 in *in vivo* ALD model. Immunofluorescent analysis confirmed these observations (Fig. 6E). As shown in Fig. 6F, the co-IP results indicated that HSD11B1 also interacted with its downstream protein IDI1. In addition, the dual-luciferase reporter assay demonstrated that *Srebp2* positively regulates *Idi1* expression (Fig. 6G). In *in vivo* experiments, it was confirmed that in normal liver tissue, the expression of *Srebp2* and *Idi1* followed a similar trend in response to overexpression or knockdown of *Hsd11b1* (Fig. 6H and 6I). Knockdown of *Srebp2* resulted in a decrease in *Idi1* expression, without affecting *Hsd11b1* levels. Conversely, neither knockdown nor overexpression of *Idi1* influenced the expression of the upstream genes *Hsd11b1* or *Srebp2*. In liver tissue from ALD, the expression changes of upstream and downstream genes upon overexpression or knockdown of *Hsd11b1* or *Idi1* mirrored those seen in normal liver tissue. However, *Srebp2* knockdown induced a negative feedback effect, reducing *Hsd11b1* expression, which was not observed in normal liver tissue (Fig. 6J and 6K).

**Fig. 6.**
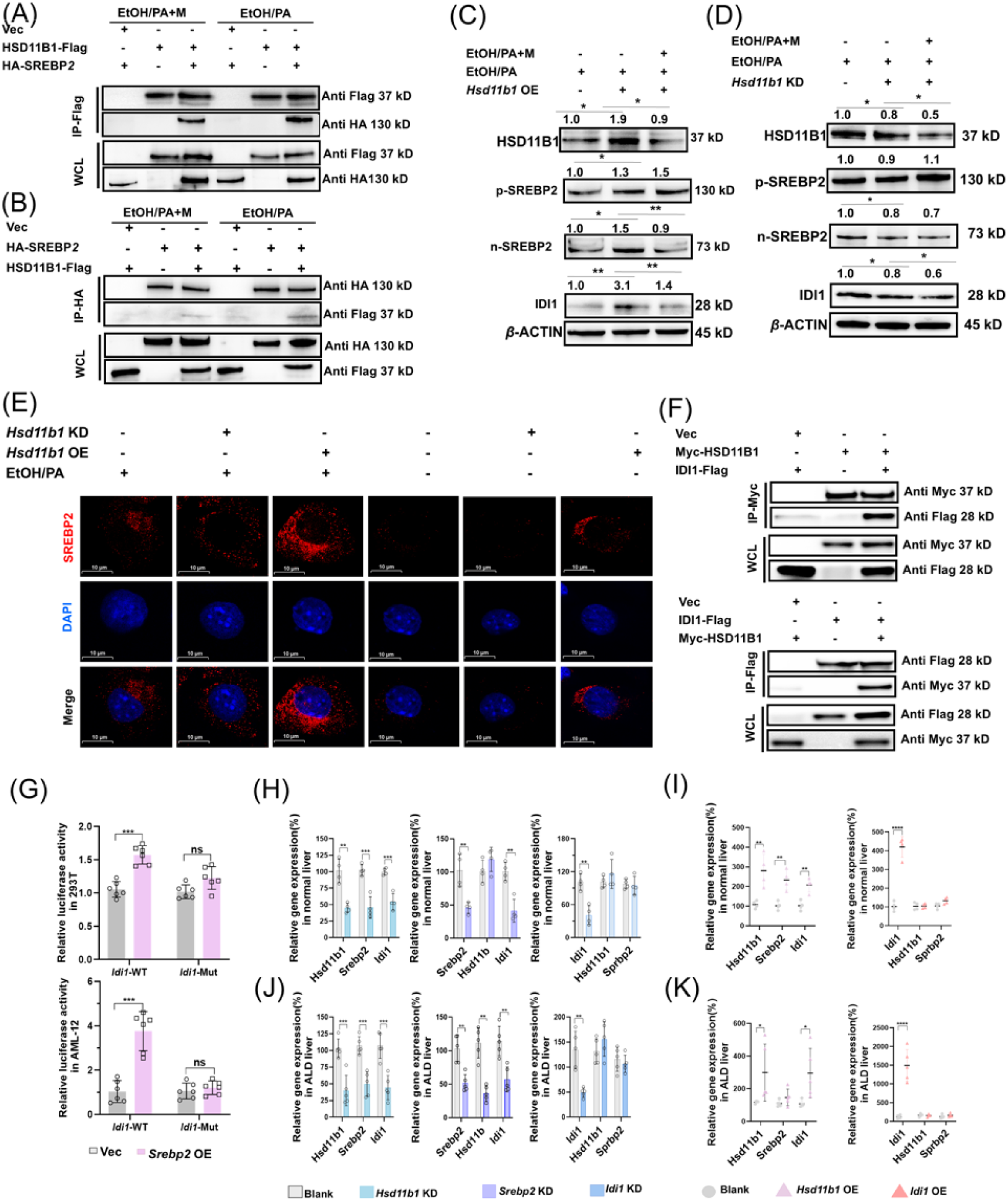
MgIG exerts its protective effect via the HSD11B1-SREBP2-IDI1 axis in ALD model. (A, B) Co-IP was used to verify the direct binding between HSD11B1 and SREBP2 with or without MgIG treatment. (C, D) Alterations in protein levels of HSD11B1, p-SREBP2, n-SREBP2, and IDI1 in AML-12 cells induced by EtOH/PA, following knockdown or overexpression of *Hsd11b1*, with and without MgIG treatment. Numbers above the lanes indicate the mean relative density normalized to the loading control for each group (n = 3). (E) Immunofluorescent staining was conducted to visualize the distribution and expression of SREBP2 (red) in EtoH/PA-induced AML-12 cells, with or without MgIG, following *Hsd11b1* knockdown or overexpression. (F) Co-IP was used to verify the direct binding between HSD11B1 and IDI1. (G) Effects of *Srebp2* on *Idi1* transcriptional regulation were measured by luciferase assays in AML-12 and 293T cell lines. Idi1-wild-type (WT) or Idi1-mutant (Mut), plasmids with WT promoter cDNA clone of *Idi1* or with mutant promoter cDNA clone plasmid; pRL-TK, an internal control reporter plasmid. (H, I) The expression levels of *Hsd11b1*, *Srebp2*, and *Idi1* in normal liver were measured following the knockdown of *Hsd11b1*, *Srebp2*, and *Idi1*, or the overexpression of *Hsd11b1* and *Idi1*, respectively (n = 4). (J, K) The expression levels of *Hsd11b1*, *Srebp2*, and *Idi1* in ALD liver were measured following the knockdown of *Hsd11b1*, *Srebp2*, and *Idi1*, or the overexpression of *Hsd11b1* and *Idi1*, respectively (n = 5). Data are expressed as mean ± SD. \**P* < 0.05, \*\**P* < 0.01, \*\*\**P* < 0.001. \*\*\*\**P* < 0.0001. Scale bar: 10 μm.

To further investigate the roles of HSD11B1, SREBP2, and IDI1 in the *in vivo* ALD model, we used AAV8-mediated shRNA delivery to knock down their expression or employed overexpression plasmids to increase their expression specifically in hepatocytes. As shown in Fig. S6A-6C, knockdown or overexpression of *Hsd11b1* or *Idi1* in mice fed a normal control diet did not result in significant changes in liver histology, lipid accumulation, or serum ALT and AST levels (Fig. S7). However, in the ALD model, knockdown of *Hsd11b1*, *Srebp2*, or *Idi1* attenuated ethanol-induced liver injury, as evidenced by improved histological NAS scores, reduced steatosis, and decreased inflammation. MgIG treatment further enhanced these beneficial effects. Conversely, overexpression of *Hsd11b1* or *Idi1* abolished the therapeutic effects of MgIG on ethanol-induced liver injury (Fig. 7A-7F and S8). Interestingly, modulation of *Idi1* expression did not affect the expression of *Hsd11b1* or *Srebp2*, suggesting that IDI1 acts downstream of HSD11B1 and SREBP2. In contrast, modulation of *Hsd11b1* expression altered the levels of n-SREBP2 and IDI1 (Fig. S8E-H), confirming that HSD11B1 regulates the SREBP2-IDI1 pathway. Collectively, these data indicate that MgIG protects against ALD, at least in part, by modulating the HSD11B1-SREBP2-IDI1 axis.

**Fig. 7.**
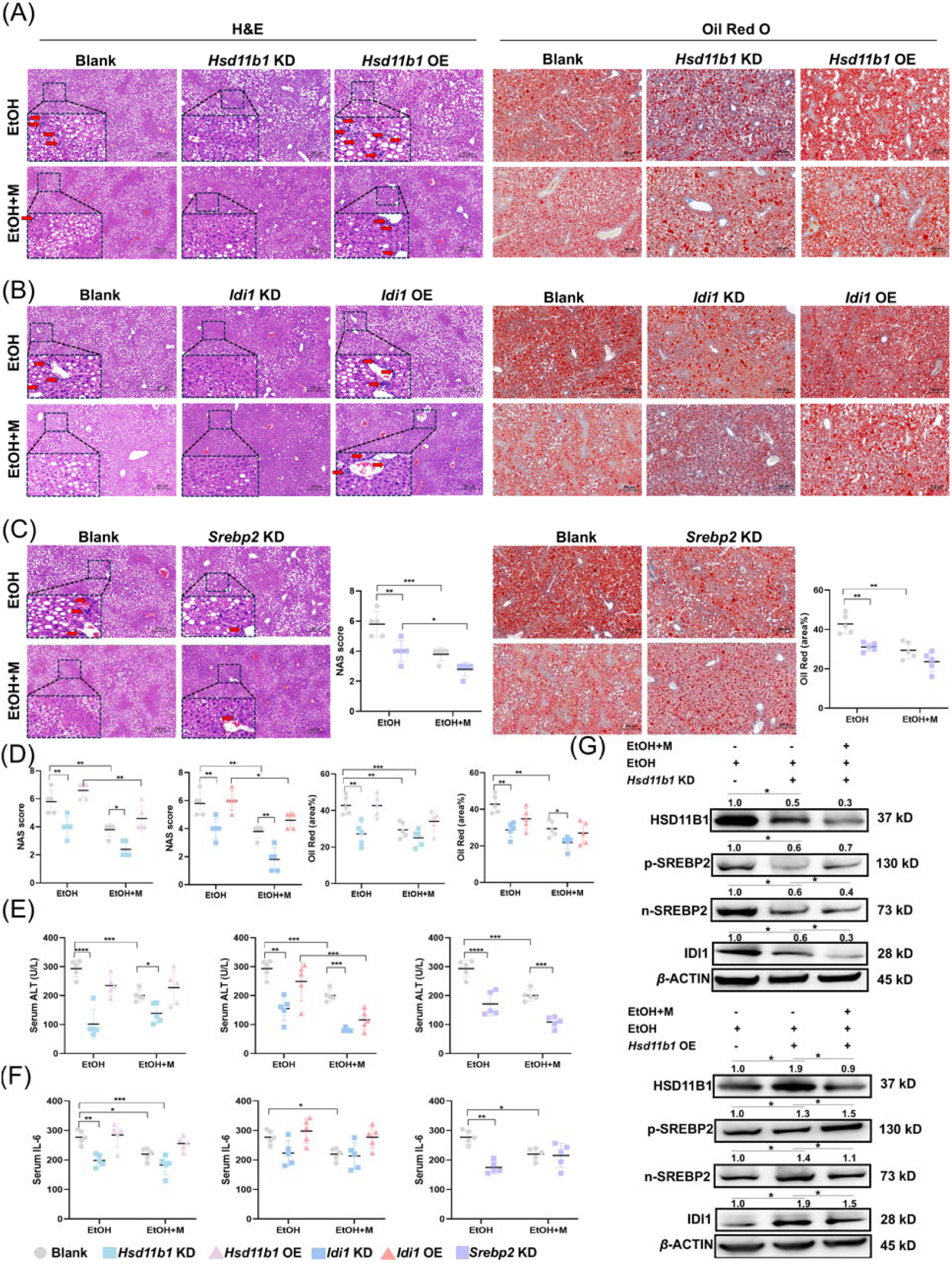
MgIG alleviates ALD-induced liver injury via the HSD11B1-SREBP2-IDI1 axis. (A-D) Representative liver H&E and Oil Red O staining results from ALD mice with *Hsd11b1*, *Srebp2*, or *Idi1* knockdown or overexpression, with and/or without MgIG co-treatment. Changes in quantitative NAS (NAFLD activity score) and Oil Red staining (area %) were calculated and analyzed (n = 5). (E-F) Changes of serum ALT and TNF-α from ALD mice with *Hsd11b1*, *Srebp2*, or *Idi1* knockdown or overexpression, with and/or without MgIG co-treatment (n = 5). (G) Alterations in protein levels of HSD11B1, p-SREBP2, n-SREBP 2, and Idi1 in ALD mice, following knockdown or overexpression of *Hsd11b1*, with and without MgIG co-treatment. Numbers above the lanes indicate the mean relative density normalized to the loading control for each group (n = 3). Data are expressed as mean ± SD. \**P* < 0.05, \*\**P* < 0.01, \*\*\**P* < 0.001, \*\*\*\**P* < 0.0001. Scale bar: 50 μm.

## Discussion

While previous research using a zebrafish model demonstrated that MgIG alleviates alcohol-induced liver injury by modulating lipid metabolism-related gene expression ^18^, the precise molecular mechanisms and direct targets of MgIG remained unclear. This lack of mechanistic understanding hindered the development of optimized therapeutic strategies for ALD. In this study, we provide compelling evidence that MgIG effectively ameliorates key features of ALD, including hepatic steatosis, inflammation, and apoptosis, in both *in vivo* and *in vitro* models. Our findings shed light on the complex interplay of molecular pathways involved in ALD pathogenesis and highlight the therapeutic potential of targeting the HSD11B1-SREBP2-IDI1 axis.

Our RNA sequencing analysis identified IDI1 as a key player in ALD pathogenesis and a critical mediator of MgIG’s hepatoprotective effects. IDI1, an enzyme in the mevalonate pathway, is involved in the biosynthesis of isoprenoids, which are essential for various cellular processes, including cholesterol synthesis. Dysregulation of cholesterol metabolism is a hallmark of ALD, contributing to the accumulation of lipids in the liver. Our findings demonstrate that *Idi1* expression is significantly upregulated in the livers of mice subjected to the NIAAA model of ALD, suggesting that increased IDI1 activity may contribute to the development of hepatic steatosis. Genetic manipulation of *Idi1* expression further confirmed its role in ALD. Specifically, *Idi1* knockdown protected against alcohol-induced liver injury, an effect further enhanced by MgIG treatment. This suggests that reducing IDI1 activity can ameliorate the detrimental effects of alcohol on the liver and that MgIG may exert its therapeutic benefits, at least in part, by suppressing IDI1. Conversely, *Idi1* overexpression abolished the therapeutic benefits of MgIG, both *in vivo* and *in vitro*, highlighting the importance of IDI1 in mediating MgIG’s effects. These findings are consistent with recent reports highlighting the role of IDI1 in hepatic cholesterol accumulation and lipid toxicity ^40^ ^41^ and its involvement in hepatocellular carcinoma progression via inhibition of the cGAS-STING pathway ^9^. Furthermore, our study identified SREBP2, a master regulator of lipid metabolism, as an upstream regulator of IDI1. SREBP2 belongs to a family of transcription factors that control the expression of genes involved in cholesterol and fatty acid biosynthesis. SREBP2 has been implicated in various aspects of liver pathology, including NLRP3 inflammasome activation ^42^, lipid accumulation ^43^, liver cancer progression ^44^, with its target protein IDI1 upregulated in parallel ^30^. Our findings demonstrate that both SREBP2 and IDI1 are upregulated in response to EtOH/PA treatment and that MgIG effectively inhibits their expression, consistent with the notion that SREBP2 contributes to the development of MASLD ^45^. The identification of SREBP2 as a key regulator of IDI1 in the context of ALD provides a mechanistic link between lipid metabolism and the pathogenesis of this disease.

To identify the direct molecular target of MgIG, we employed a combination of molecular docking, MD, MST analysis, and genetical manipulation studies. These approaches converged on HSD11B1, a key enzyme in glucocorticoid metabolism, as a high-affinity binding partner of MgIG. HSD11B1 catalyzes the conversion of inactive cortisone to active cortisol in the liver, thereby increasing intracellular glucocorticoid levels. Glucocorticoids are steroid hormones that play a critical role in regulating glucose and lipid metabolism, and their dysregulation has been implicated in the development of metabolic disorders, including type 2 diabetes ^46^. As a potential inhibitor of HSD11B1 ^47^, GL can improve patients’ insulin and lipid levels ^48^. In addition, the increased expression of *Hsd11b1* enhances the enzymatic reducing activity, promoting a fat-forming phenotype in cell models through elevated cortisol levels ^49,50^. Meanwhile, *Hsd11b1* deficiency prevents hepatic steatosis in high-fat diets-fed mice ^51^. Our findings demonstrate that MgIG directly binds to the Lys187 of HSD11B1 and that modulation of *Hsd11b1* expression in hepatocytes significantly influences the development of ALD.

While some studies have suggested a potential pro-fibrotic role for HSD11B1 inhibition ^8^, our findings, along with those of other recent studies ^7,51,52^, support a protective role for *Hsd11b1* knockdown in the context of alcoholic liver injury. These discrepancies may be attributed to differences in experimental models and the specific cell types targeted. For instance, the study by Zou *et al* ^8^ examined the effects of global *Hsd11b1* deficiency, which could trigger compensatory activation of the hypothalamic-pituitary-adrenal axis or paracrine regulatory mechanisms, potentially affecting the pro-fibrotic response. In contrast, our study focused on hepatocyte-specific *Hsd11b1* knockdown, which may avoid these confounding factors. Importantly, our study provides evidence for a novel HSD11B1-SREBP2-IDI1 axis in the pathogenesis of ALD. We demonstrate that HSD11B1 regulates the processing and nuclear translocation of SREBP2, which in turn controls the expression of IDI1. Notably, this is consistent with previous studies showing that increased SREBP2 expression and nuclear translocation activate the NLRP3 inflammasome and the mevalonate pathway, thereby promoting inflammation and tumor progression ^53,54^. This regulatory axis appears to be a key target of MgIG’s therapeutic effects. By directly binding to HSD11B1, MgIG may inhibit its activity, leading to reduced SREBP2 activation and subsequent downregulation of IDI1. This, in turn, could contribute to the observed improvements in hepatic steatosis, inflammation, and apoptosis ^55–57^.

This study provides mechanistic evidence that MgIG directly targets HSD11B1 and modulates the HSD11B1–SREBP2–IDI1 axis to attenuate ALD. However, several limitations should be acknowledged. First, our findings are derived from preclinical mouse models and hepatocyte systems, and pharmacokinetics, biodistribution, and long-term safety of MgIG remain to be comprehensively evaluated in larger and more diverse populations. Second, although molecular docking and thermophoresis confirmed direct binding of MgIG to HSD11B1, high-resolution structural studies such as X-ray crystallography or cryo-electron microscopy are needed to delineate the precise binding interface and conformational changes. Third, other signaling pathways potentially influenced by MgIG have not been systematically explored and may contribute to its therapeutic effects. Future work should focus on validating these findings in human tissues, optimizing MgIG dosing strategies, and conducting multicenter clinical trials to assess its translational potential and establish HSD11B1 as a druggable target for precision therapy in ALD.

## Supporting information

Body weight

## Acknowledgements

JX is supported by National Natural Science Foundation of China (U23A20401) and Science and Technology Projects in Guangzhou (202201020066). YL is supported by National Natural Science Foundation of China (82372768). HZ is supported by Medical Research Fund of The Six Affiliated Hospital of Jinan University.

## Authorcontributions

**Jia Xiao**: Supervision, Conceptualization, Funding acquisition, Writing– review & editing, Data curation. **Yan Li**: Funding acquisition, Conceptualization, Data curation, Writing– review & editing. **Lu Xiao**: Writing original draft, Methodology, Data curation, Visualization. **Lu Li**: Writing original draft, Methodology, Data curation, Visualization. **Shasha Wu**: Data curation, Methodology. **Zhaoyi Che**: Data curation, Methodology. **Yuyang Du**: Data curation, Methodology. **Jingyi Zheng**: Data curation, Methodology. **Jingsong Yan**: Data curation. **Hua Wang:** Wrote and revised the manuscript. **Hong Zhang:** Funding acquisition, Wrote and revised the manuscript.

## Data availability

All data is available in the manuscript or the supplementary materials. RNA-seq data from this study is available from: GSE284311. All materials generated as part of this study will be made available upon request to the corresponding authors.

## Declaration ofinterests

The authors declare no competing interests.

## Abbreviations

ALD: alcohol-associated liver disease
ALT: alanine aminotransferase
AST: aspartate aminotransferase
DEG: differentially expressed gene
DILI: drug-induced liver injury
GL: generation glycyrrhizin
H&E: hematoxylin and eosin
HSD11B1: hydroxysteroid 11-beta dehydrogenase 1
IDI1: isopentenyl diphosphate delta isomerase 1
IL-6: interleukin-6
KEGG: Kyoto Encyclopedia of Genes and Genomes
LDH: lactate dehydrogenase
MASLD: metabolic dysfunction-associated steatotic liver disease
MgIG: magnesium isoglycyrrhizinate
MST: microscale thermophoresis
NAS: nonalcoholic fatty liver disease activity score
NIAAA: National Institute on Alcohol Abuse and Alcoholism
n-Srebp2: nuclear SREBP2
PA: palmitic acid
PBS: phosphate-buffered saline
p-Srebp2: precursor SREBP2
SREBP2: sterol regulatory element binding protein 2
TC: total cholesterol
TG: triglyceride
TNF-α: tumor necrosis factor-alpha
ULM: univariate linear model.

## Supporting figures

**Fig. S1.**
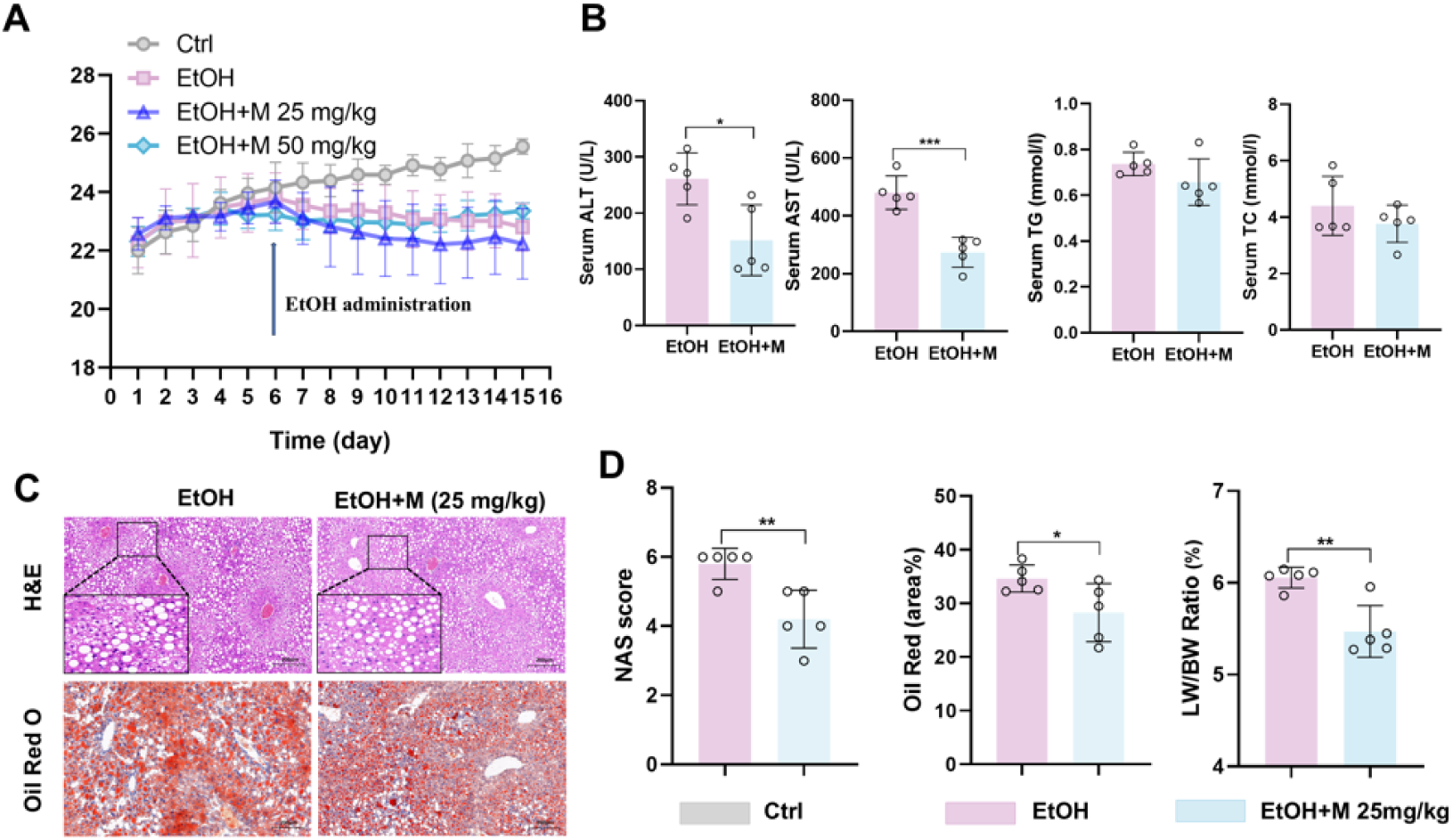
The impact of 25 mg/kg MgIG on liver injury parameters in mice. (A) Changes in body weight of mice with or without dietary and alcohol induction, and with or without MgIG treatment (n = 5). (B) Changes of NAS (NAFLD activity score), Oil Red O quantification (area%), with ALD and/or MgIG treatment (25 mg/kg; n = 5). (C) Representative liver H&E and Oil Red O staining results from mice with ALD and/or MgIG (25 mg/kg) treatment. (D) Changes of serum biochemical parameters (ALT, AST and TG) from mice with ALD and/or MgIG treatment (25 mg/kg; n = 5). *P < 0.05, **P < 0.01, ***P < 0.001.

**Fig. S2.**
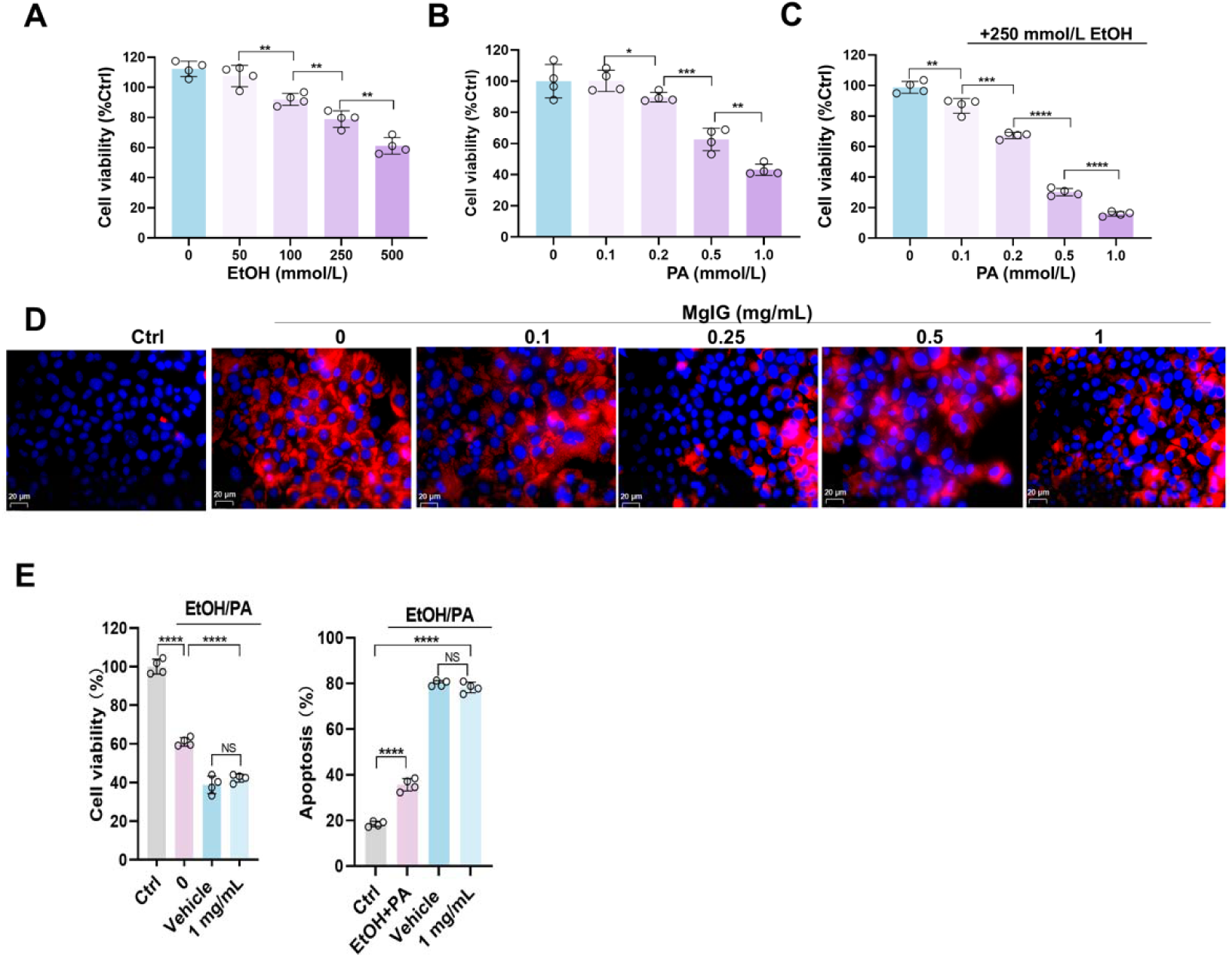
Optimization tests of the concomitant treatment doses of ethanol and palmitic acid (PA) in mouse normal hepatocyte AML-12 cell line, to partially induce alcohol-associated liver disease (ALD) phenotypes. (A) Changes of cell viability after a series of doses of ethanol PA treatment (n = 4). (B) Changes of cell viability after a series of doses of ethanol treatment (n = 4). (C) Changes of cell viability after different doses of PA and 250 mmol/L EtOH concurrent treatments (n = 4). (D) Representative low-magnification images of Nile Red staining in AML-12 cells treated with ethanol/PA, with or without co-treatment with different doses of MgIG. (E) Viability/toxicity of the 1.0 mg/mL group with vehicle control. *P < 0.05, **P < 0.01, ***P < 0.001, ****P <0.0001. Scale bar: 20 μm.

**Fig. S3.**
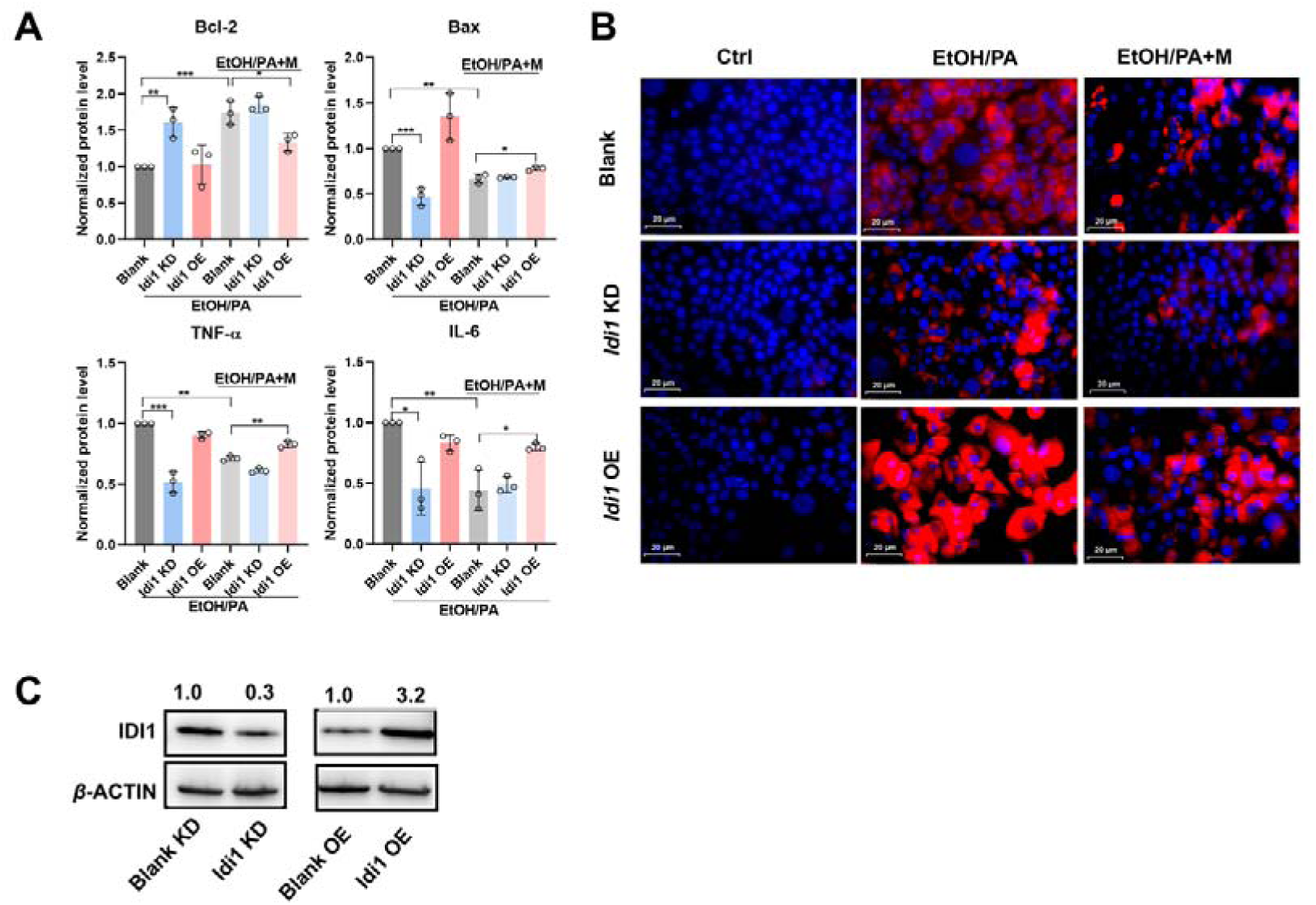
IDI1 is involved in MgIG-mediated hepatocyte protection against ethanol. (A) Densitometric analysis of inflammatory (TNF-_α_ and IL-6) and apoptotic (Bax and Bcl-2) factors in AML-12 cells with *Idi1* knockdown or overexpression following EtOH/PA treatment, with or without MgIG co-treatment. Data are expressed as mean ± SD (n = 3). (B) Representative low-magnification images of Nile Red staining in AML-12 cells with *Idi1* knockdown or overexpression, treated with ethanol/PA, with or without co-treatment with different doses of MgIG. (C) IDI1 overexpression and knockdown efficiency (Western blot). *P < 0.05, **P < 0.01, ***P < 0.001, ****P <0.0001. Scale bar: 20 μm.

**Fig. S4.**
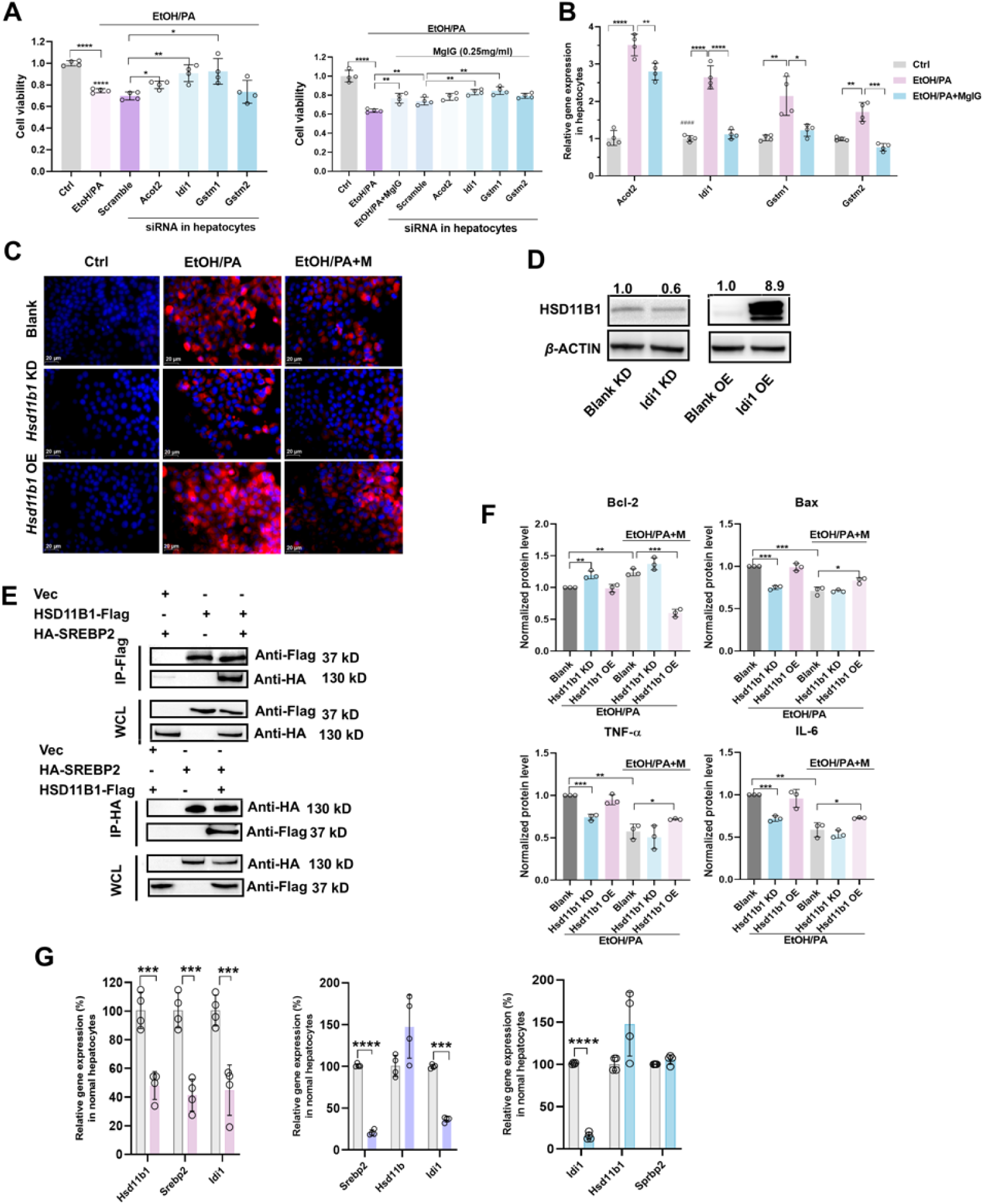
*Hsd11b1/Idi1* is involved in ethanol/palmitic acid (PA) concomitantly induced AML-12 hepatocyte damage and MgIG protection. (A) Changes of cell viability ratio of AML-12 cells treated with ethanol/PA in the absence or presence of 0.25 mg/mL MgIG, and knock-down of specified genes (n = 4). (B) Quantitative PCR validation of RNA-seq exhibited top differentiated genes in AML-12 cells treated with ethanol/PA in the absence or presence of 0.25 mg/mL MgIG (n = 4). (C) Representative low-magnification images of Nile Red staining in AML-12 cells with *Hsd11b1* knockdown or overexpression, treated with ethanol/PA, with or without co-treatment with different doses of MgIG. (D) HSD11B1 overexpression and knockdown efficiency (Western blot). (E) Western Blot after Co-IP was used to verify the binding of HSD11B1 with SREBP2. (F) The expression levels of *Hsd11b1*, *Srebp2*, and *Idi1* in hepatocytes following the knockdown of HSD11B1, SREBP2, and IDI1, respectively (n = 4). (F) Densitometric analysis of inflammatory (TNF-_α_ and IL-6) and apoptotic (Bax and Bcl-2) factors in AML-12 cells with *Hsd11b1* knockdown or overexpression following EtOH/PA treatment, with or without MgIG co-treatment. Data are expressed as mean ± SD (n = 3). (G) The expression levels of *Hsd11b1*, *Srebp2*, and *Idi1* in hepatocytes following the knockdown of HSD11B1, SREBP2, and IDI1, respectively (n = 4). *P < 0.05, **P < 0.01, ***P < 0.001, ****P <0.0001.

**Fig. S5.**
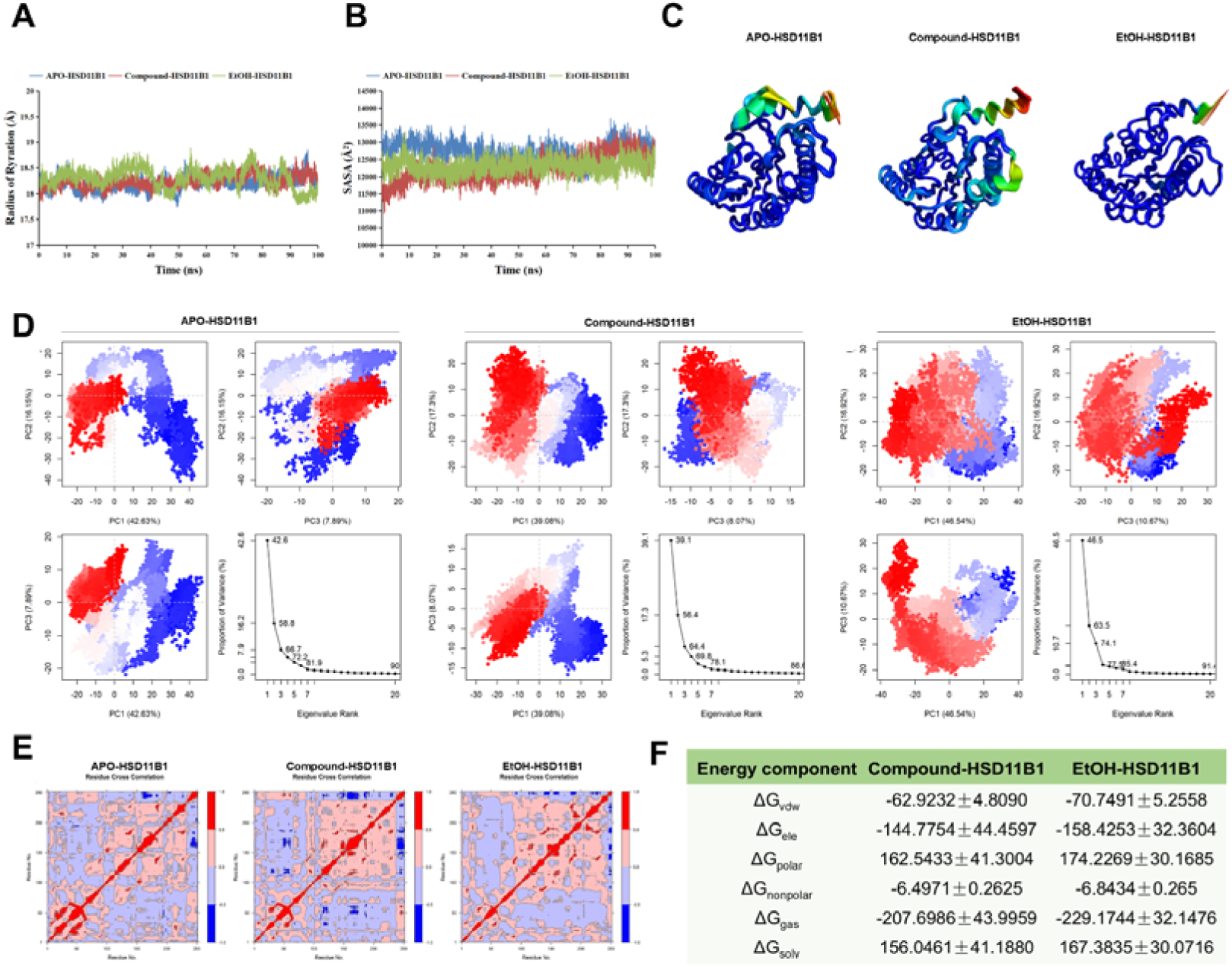
MgIG directly binds to HSD11B1. (A) Analysis of radius of gyration of three systems. (B) Solvent accessible surface area (SASA) analysis of the three systems. (C-D) Principal component analysis (PCA) of the three systems. (E) Cross-correlation analysis of three systems. (F) Binding free energy of Compound-HSD11B1 and EtOH-HSD11B1. APO-HSD11B1 represents unbound HSD11B1 in a physiological saline system, Compound-HSD11B1 represents HSD11B1 bound to MgIG in a physiological saline system, and EtOH-HSD11B1 represents HSD11B1 bound to MgIG in a 0.1 mg/mL ethanol solvent system.

**Fig. S6.**
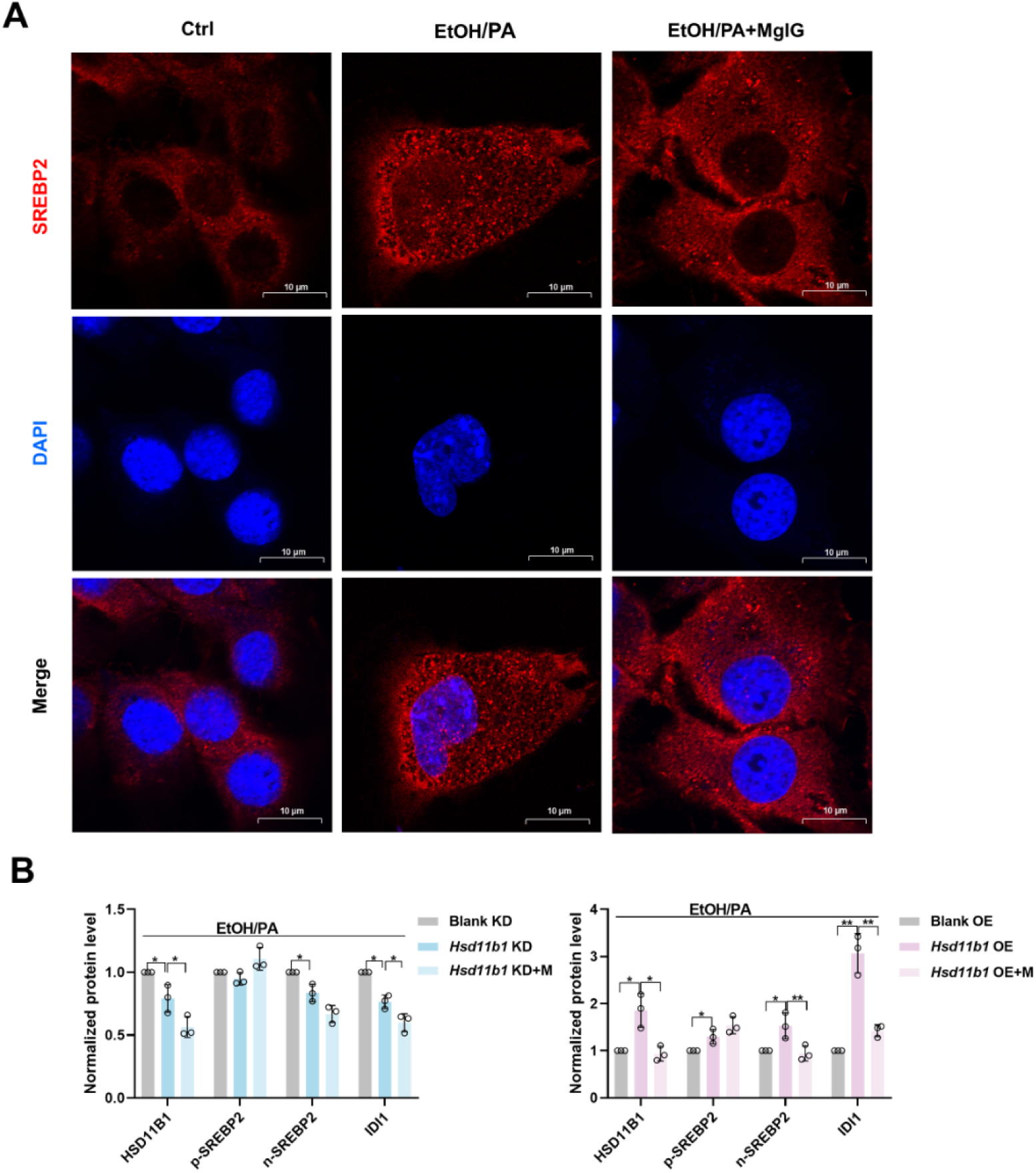
(A) Immunofluorescent staining was conducted to visualize the distribution and expression of SREBP2 (red) in EtOH/PA-induced AML-12 cells, with or without 0.25 mg/mL MgIG. (B) Densitometric analysis of HSD11B1, SREBP2 and IDI1 protein levels (normalized to β-actin) in AML-12 cells with *Hsd11b1* knockdown or overexpression following EtOH/PA treatment, with or without MgIG co-treatment. Data are expressed as mean ± SD (n = 3). Scale bar: 7.5 μm. EtOH, ethanol. PA, palmitic acid.

**Fig. S7.**
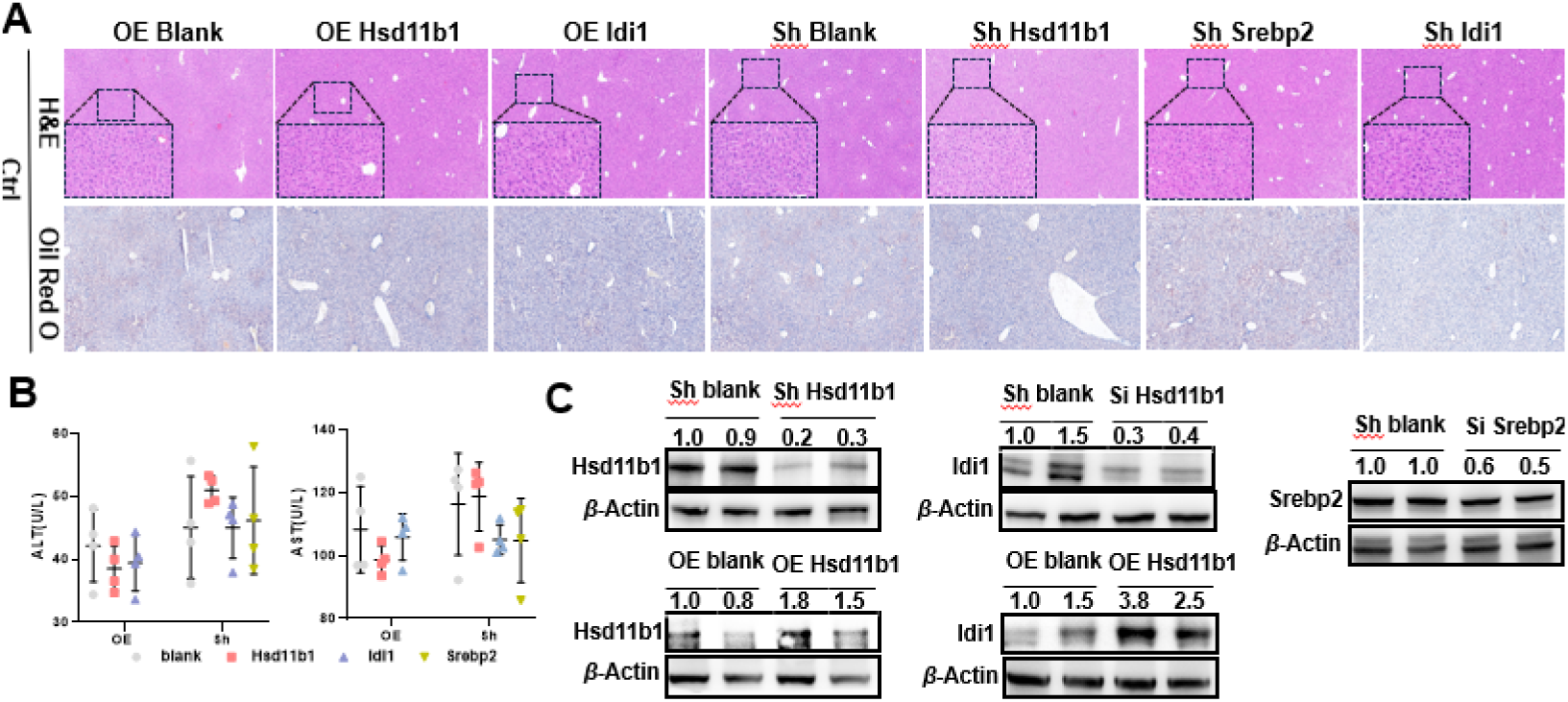
Knockdown and overexpression of *Hsd11b1/Srebp2/Idi1* do not affect liver function in mice fed a normal diet. (A) Representative liver H&E and Oil Red O staining results of normal diet mice with knockdown of *Hsd11b1/Srebp2/Idi1* or overexpression of *Hsd11b1/Idi1*. (B) Changes in serum biochemical parameters (ALT and AST) of normal diet mice, with or without *Hsd11b/Serep2/Idi1* knockdown or *Hsd11b/Idi1* overexpression (n = 4). (C) Validation of knockdown efficiency of *Hsd11b1/Srebp2/Idi1* and overexpression efficiency of *Hsd11b1/Idi1* in mice fed a normal diet by using Western blot analysis. Scale bar: 50 μm. Si, siRNA knockdown. OE, overexpression.

**Fig. S8.**
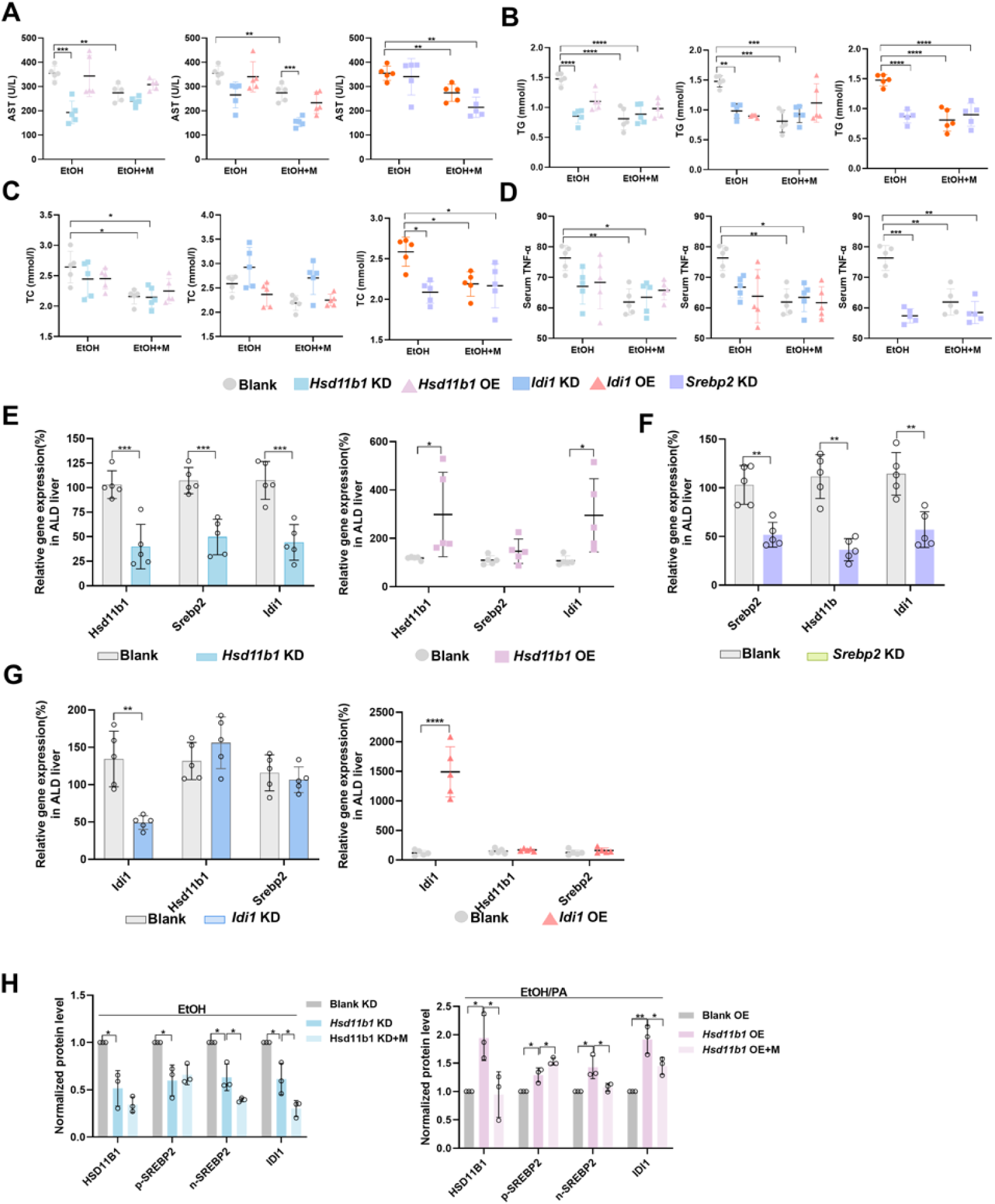
MgIG exerts its protective effect via the HSD11B1-SREBP2-IDI1 axis in mice ALD modes. (A-D) Changes of serum AST, TG, TC and TNF-α from ALD mice with *Hsd11b1, Srebp2*, or *Idi1* knockdown or overexpression, with and/or without MgIG treatment (n = 5). (E) Changes in downstream gene expression after *Hsd11b1* knockdown and overexpression (n = 5). (F) Changes in the expression of upstream and downstream genes after *Srebp2* knockdown (n = 5). (G) Changes in upstream gene expression after *Idi1* knockdown and overexpression (n = 5). (H) Densitometric analysis of HSD11B1, SREBP2, and IDI1 protein levels (normalized to β-actin) in livers from ethanol-induced liver injury mice with *Hsd11b1* knockdown or overexpression, with or without MgIG co-treatment. Data are expressed as mean ± SD (n = 3). *P < 0.05, **P < 0.01, ***P < 0.001, ****P <0.0001. KD, knockdown. OE, overexpression.

## 2. Supporting dataset

### Uncropped Western blot data

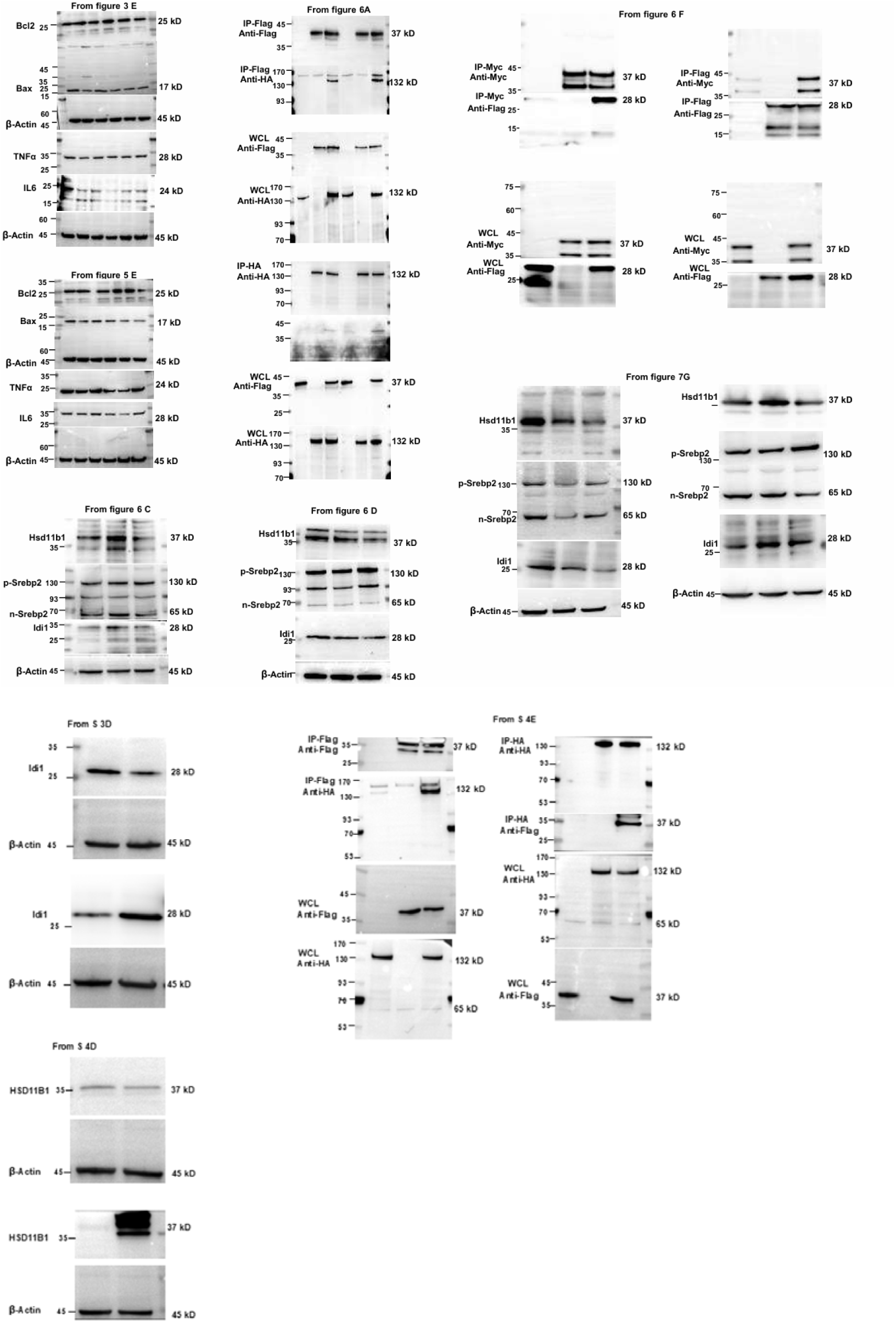

## 3. Supporting tables

**Table S1.**
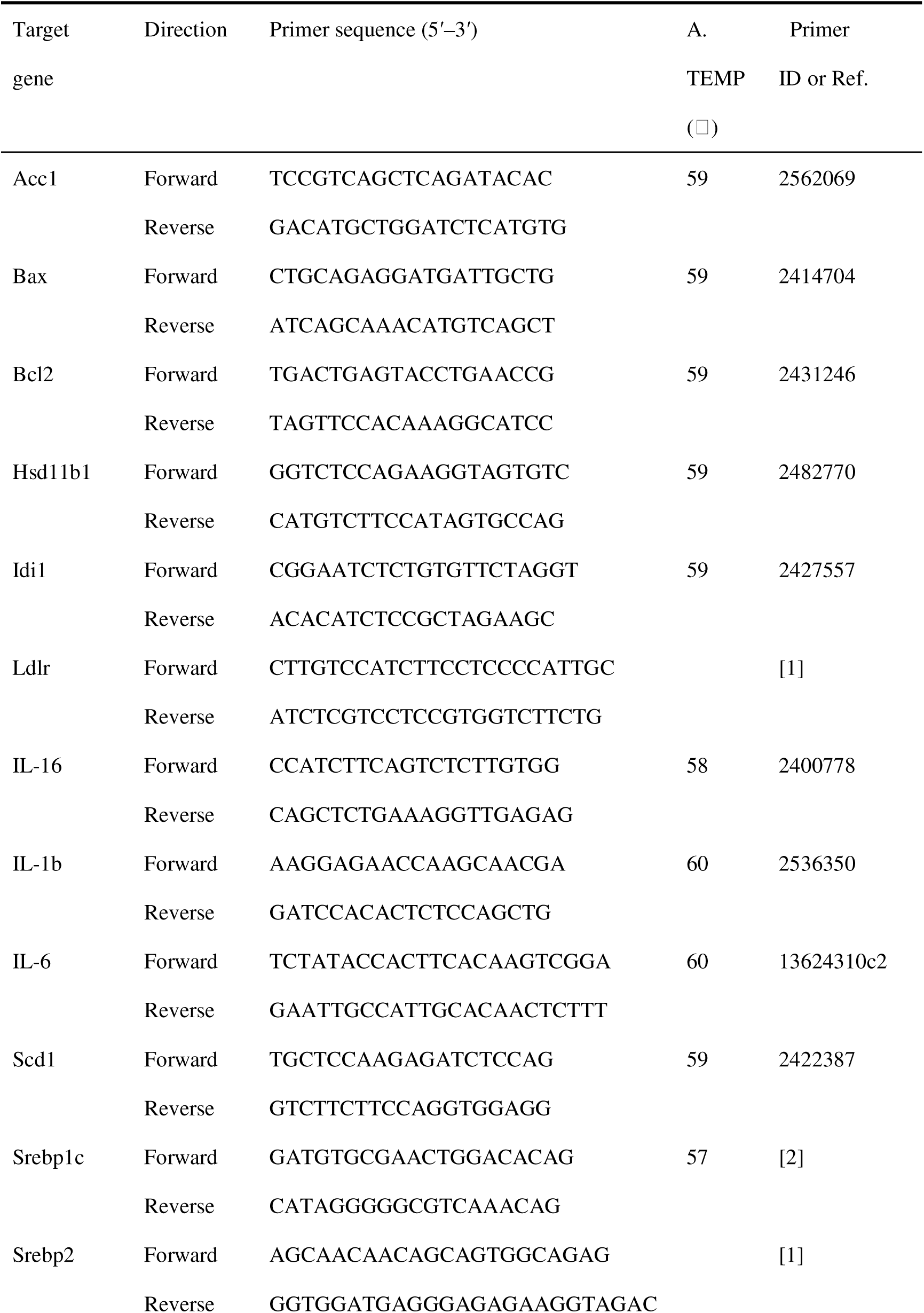
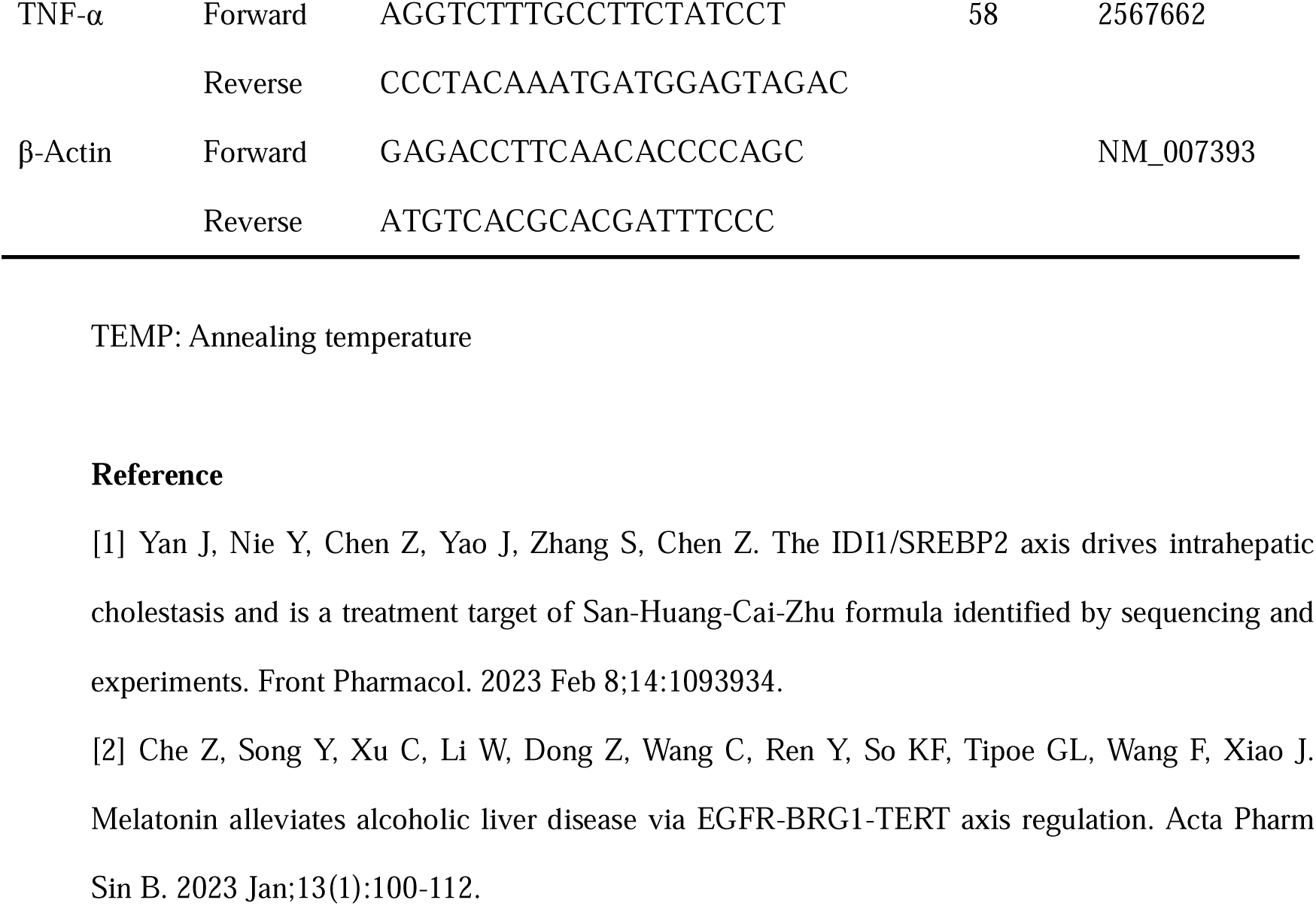
Primer sequences information for quantitative real-time PCR assay.

## REFERENCES

1. Aberg, F., Jiang, Z.G., Cortez-Pinto, H., and Mannisto, V. (2024). Alcohol-associated liver disease-Global epidemiology. Hepatology. 10.1097/HEP.0000000000000899.

2. Gan, C., Yuan, Y., Shen, H., Gao, J., Kong, X., Che, Z., Guo, Y., Wang, H., Dong, E., and Xiao, J. (2025). Liver diseases: epidemiology, causes, trends and predictions. Signal Transduct Target Ther 10, 33. 10.1038/s41392-024-02072-z.

3. Devarbhavi, H., Asrani, S.K., Arab, J.P., Nartey, Y.A., Pose, E., and Kamath, P.S. (2023). Global burden of liver disease: 2023 update. J Hepatol 79, 516–537. 10.1016/j.jhep.2023.03.017.

4. Díaz, L.A., Arab, J.P., Louvet, A., Bataller, R., and Arrese, M. (2023). The intersection between alcohol-related liver disease and nonalcoholic fatty liver disease. Nat Rev Gastroenterol Hepatol 20, 764–783. 10.1038/s41575-023-00822-y.

5. Wu, X., Fan, X., Miyata, T., Kim, A., Cajigas-Du Ross, C.K., Ray, S., Huang, E., Taiwo, M., Arya, R., Wu, J., et al. (2023). Recent Advances in Understanding of Pathogenesis of Alcohol-Associated Liver Disease. Annu Rev Pathol 18, 411–438. 10.1146/annurev-pathmechdis-031521-030435.

6. Gao, H., Jiang, Y., Zeng, G., Huda, N., Thoudam, T., Yang, Z., Liangpunsakul, S., and Ma, J. (2024). Cell-to-cell and organ-to-organ crosstalk in the pathogenesis of alcohol-associated liver disease. eGastroenterology 2. 10.1136/egastro-2024-100104.

7. Li, C., Li, M., Sheng, W., Zhou, W., Zhang, Z., Ji, G., and Zhang, L. (2024). High dietary Fructose Drives Metabolic Dysfunction-Associated Steatotic Liver Disease via Activating ubiquitin-specific peptidase 2/11beta-hydroxysteroid dehydrogenase type 1 Pathway in Mice. Int J Biol Sci 20, 3480–3496. 10.7150/ijbs.97309.

8. Zou, X., Ramachandran, P., Kendall, T.J., Pellicoro, A., Dora, E., Aucott, R.L., Manwani, K., Man, T.Y., Chapman, K.E., Henderson, N.C. et al. (2018). 11Beta-hydroxysteroid dehydrogenase-1 deficiency or inhibition enhances hepatic myofibroblast activation in murine liver fibrosis. Hepatology 67, 2167–2181. 10.1002/hep.29734.

9. Fu, L., Ding, H., Bai, Y., Cheng, L., Hu, S., and Guo, Q. (2024). IDI1 inhibits the cGAS-Sting signaling pathway in hepatocellular carcinoma. Heliyon 10. 10.1016/j.heliyon.2024.e27205.

10. Yan, J.B., Nie, Y.M., Chen, Z., Yao, J.M., Zhang, S., and Chen, Z.Y. (2023). The IDI1/SREBP2 axis drives intrahepatic cholestasis and is a treatment target of San-Huang-Cai-Zhu formula identified by sequencing and experiments. Front Pharmacol 14. https://doi.org/ARTN 109393410.3389/fphar.2023.1093934.

11. Fan, X., Wang, H., Wang, W., Shen, J., and Wang, Z. (2024). Exercise training alleviates cholesterol and lipid accumulation in mice with non-alcoholic steatohepatitis: Reduction of KMT2D-mediated histone methylation of IDI1. Exp Cell Res 442, 114265. 10.1016/j.yexcr.2024.114265.

12. Wang, L., Yang, R., Yuan, B., Liu, Y., and Liu, C. (2015). The antiviral and antimicrobial activities of licorice, a widely-used Chinese herb. Acta Pharm Sin B 5, 310–315. 10.1016/j.apsb.2015.05.005.

13. Ma, D., Zhang, J., Zhang, Y., Zhang, X., Han, X., Song, T., Zhang, Y., and Chu, L. (2018). Inhibition of myocardial hypertrophy by magnesium isoglycyrrhizinate through the TLR4/NF-κB signaling pathway in mice. Int Immunopharmacol 55, 237–244. 10.1016/j.intimp.2017.12.019.

14. Liu, M., Zheng, B., Liu, P., Zhang, J., Chu, X., Dong, C., Shi, J., Liang, Y., Chu, L., Liu, Y. et al. (2021). Exploration of the hepatoprotective effect and mechanism of magnesium isoglycyrrhizinate in mice with arsenic trioxide-induced acute liver injury. Mol Med Rep 23. 10.3892/mmr.2021.12077.

15. Lai, X., Zhou, H., Wan, Y., Kuang, J., Yang, Y., Mai, L., Chen, Y., and Liu, B. (2024). Magnesium isoglycyrrhizinate attenuates nonalcoholic fatty liver disease by strengthening intestinal mucosal barrier. Int Immunopharmacol 128, 111429. 10.1016/j.intimp.2023.111429.

16. Yang, Y.-Z., Zhao, X.-J., Xu, H.-J., Wang, S.-C., Pan, Y., Wang, S.-J., Xu, Q., Jiao, R.-Q., Gu, H.-M., and Kong, L.-D. (2019). Magnesium isoglycyrrhizinate ameliorates high fructose-induced liver fibrosis in rat by increasing miR-375-3p to suppress JAK2/STAT3 pathway and TGF-β1/Smad signaling. Acta Pharmacol Sin 40, 879–894. 10.1038/s41401-018-0194-4.

17. Lu, C., Xu, W., Shao, J., Zhang, F., Chen, A., and Zheng, S. (2017). Blockade of hedgehog pathway is required for the protective effects of magnesium isoglycyrrhizinate against ethanol-induced hepatocyte steatosis and apoptosis. IUBMB Life 69, 540–552. 10.1002/iub.1639.

18. Dai, W., Wang, K., Zhen, X., Huang, Z., and Liu, L. (2022). Magnesium isoglycyrrhizinate attenuates acute alcohol-induced hepatic steatosis in a zebrafish model by regulating lipid metabolism and ER stress. Nutr Metab (Lond) 19, 23. 10.1186/s12986-022-00655-7.

19. National Workshop on Fatty, L., Alcoholic Liver Disease, C.S.o.H.C.M.A., and Fatty Liver Expert Committee, C.M.D.A. (2018). [Guidelines of prevention and treatment for alcoholic liver disease: a 2018 update]. Zhonghua Gan Zang Bing Za Zhi 26, 188–194. 10.3760/cma.j.issn.1007-3418.2018.03.007.

20. Bertola, A., Mathews, S., Ki, S.H., Wang, H., and Gao, B. (2013). Mouse model of chronic and binge ethanol feeding (the NIAAA model). Nat Protoc 8, 627–637. 10.1038/nprot.2013.032.

21. Wang, Y., Zhang, Z., Wang, X., Qi, D., Qu, A., and Wang, G. (2017). Amelioration of Ethanol-Induced Hepatitis by Magnesium Isoglycyrrhizinate through Inhibition of Neutrophil Cell Infiltration and Oxidative Damage. Mediators Inflamm 2017, 3526903. 10.1155/2017/3526903.

22. Brunt, E.M., Kleiner, D.E., Wilson, L.A., Belt, P., Neuschwander-Tetri, B.A., and Network, N.C.R. (2011). Nonalcoholic Fatty Liver Disease (NAFLD) Activity Score and the Histopathologic Diagnosis in NAFLD: Distinct Clinicopathologic Meanings. Hepatology 53, 810–820. 10.1002/hep.24127.

23. Che, Z., Song, Y., Xu, C., Li, W., Dong, Z., Wang, C., Ren, Y., So, K.-F., Tipoe, G.L., Wang, F. et al. (2023). Melatonin alleviates alcoholic liver disease via EGFR-BRG1-TERT axis regulation. Acta Pharm Sin B 13, 100–112. 10.1016/j.apsb.2022.06.015.

24. Yi, H.W., Ma, Y.X., Wang, X.N., Wang, C.F., Lu, J., Cao, W., and Wu, X.D. (2015). Ethanol promotes saturated fatty acid-induced hepatoxicity through endoplasmic reticulum (ER) stress response. Chin J Nat Med 13, 250–256. 10.1016/s1875-5364(15)30011-x.

25. Qiu, F., Zeng, R., Li, D., Ye, T., Xu, W., Wang, X., Yan, X., Li, H., and Hu, X. (2023). Establishment and bioinformatics evaluation of the ethanol combined with palmitic acid-induced mouse hepatocyte AFLD model (the Hu-Qiu Model). Heliyon 9, e19359. 10.1016/j.heliyon.2023.e19359.

26. Sharma, R.R., Rashid, H., Bhat, A.M., Sajeeda, A., Gupta, R., and Abdullah, S.T. (2023). Glabridin ameliorates intracellular events caused by palmitic acid and alcohol in mouse hepatocytes and fast food diet and alcohol -induced steatohepatitis and fibrosis in C57BL/6J mice model. Food Chem Toxicol 180, 114038. 10.1016/j.fct.2023.114038.

27. Bustin, S.A., Benes, V., Garson, J.A., Hellemans, J., Huggett, J., Kubista, M., Mueller, R., Nolan, T., Pfaffl, M.W., Shipley, G.L. et al. (2009). The MIQE guidelines: minimum information for publication of quantitative real-time PCR experiments. Clin Chem 55, 611–622. 10.1373/clinchem.2008.112797.

28. Badia, I.M.P., Velez Santiago, J., Braunger, J., Geiss, C., Dimitrov, D., Muller-Dott, S., Taus, P., Dugourd, A., Holland, C.H., Ramirez Flores, R.O. et al. (2022). decoupleR: ensemble of computational methods to infer biological activities from omics data. Bioinform Adv 2, vbac016. 10.1093/bioadv/vbac016.

29. Meijnikman, A.S., Davids, M., Herrema, H., Aydin, O., Tremaroli, V., Rios-Morales, M., Levels, H., Bruin, S., de Brauw, M., Verheij, J., et al. (2022). Microbiome-derived ethanol in nonalcoholic fatty liver disease. Nature Medicine 28, 2100–2106. 10.1038/s41591-022-02016-6.

30. Norrmen, C., Figlia, G., Lebrun-Julien, F., Pereira, J.A., Trotzmuller, M., Kofeler, H.C., Rantanen, V., Wessig, C., van Deijk, A.L., Smit, A.B., et al. (2014). mTORC1 controls PNS myelination along the mTORC1-RXRgamma-SREBP-lipid biosynthesis axis in Schwann cells. Cell Rep 9, 646–660. 10.1016/j.celrep.2014.09.001.

31. https://www.niaaa.nih.gov/alcohols-effects-health/alcohol-drinking-patterns

32. Wang, J., Wang, W., Kollman, P.A., and Case, D.A. (2006). Automatic atom type and bond type perception in molecular mechanical calculations. J Mol Graph Model 25, 247–260. 10.1016/j.jmgm.2005.12.005.

33. Tian, C., Kasavajhala, K., Belfon, K.A.A., Raguette, L., Huang, H., Migues, A.N., Bickel, J., Wang, Y., Pincay, J., Wu, Q. et al. (2019). ff19SB: Amino-Acid-Specific Protein Backbone Parameters Trained against Quantum Mechanics Energy Surfaces in Solution. Journal of Chemical Theory and Computation 16, 528–552. 10.1021/acs.jctc.9b00591.

34. Wang, J., Wolf, R.M., Caldwell, J.W., Kollman, P.A., and Case, D.A. (2004). Development and testing of a general amber force field. J Comput Chem 25, 1157–1174. 10.1002/jcc.20035.

35. Song, X., Bao, L., Feng, C., Huang, Q., Zhang, F., Gao, X., and Han, R. (2024). Accurate Prediction of Protein Structural Flexibility by Deep Learning Integrating Intricate Atomic Structures and Cryo-EM Density Information. Nat Commun 15, 5538. 10.1038/s41467-024-49858-x.

36. Zheng, N., Cai, Y., Zhang, Z., Zhou, H., Deng, Y., Du, S., Tu, M., Fang, W., and Xia, X. (2025). Tailoring industrial enzymes for thermostability and activity evolution by the machine learning-based iCASE strategy. Nat Commun 16, 604. 10.1038/s41467-025-55944-5.

37. Valdés-Tresanco, M.S., Valdés-Tresanco, M.E., Valiente, P.A., and Moreno, E. (2021). gmx_MMPBSA: A New Tool to Perform End-State Free Energy Calculations with GROMACS. J Chem Theory Comput 17, 6281–6291. 10.1021/acs.jctc.1c00645.

38. Xu, Z., Liu, D., Liu, D., Ren, X., Liu, H., Qi, G., Zhou, Y., Wu, C., Zhu, K., Zou, Z. et al. (2022). Equisetin is an anti-obesity candidate through targeting 11β-HSD1. Acta Pharm Sin B 12, 2358–2373. 10.1016/j.apsb.2022.01.006.

39. Lai, X., Zhou, H., Wan, Y., Kuang, J., Yang, Y., Mai, L., Chen, Y., and Liu, B. (2024). Magnesium isoglycyrrhizinate attenuates nonalcoholic fatty liver disease by strengthening intestinal mucosal barrier. Int Immunopharmacol 128, 111429. 10.1016/j.intimp.2023.111429.

40. Lei, Z., Rong, H., Yang, Y., Yu, S., Zhang, T., Chen, L., Nie, Y., Song, Q., Hu, Q., and Guo, J. (2022). Loperamide induces excessive accumulation of bile acids in the liver of mice with different diets. Toxicology 477, 153278. 10.1016/j.tox.2022.153278.

41. Lu, L., Hao, K., Hong, Y., Liu, J., Zhu, J., Jiang, W., Zhu, Z., Wang, G., and Peng, Y. (2021). Magnesium Isoglycyrrhizinate Reduces Hepatic Lipotoxicity through Regulating Metabolic Abnormalities. Int J Mol Sci 22. 10.3390/ijms22115884.

42. Wang, X., Wu, F.P., Huang, Y.R., Li, H.D., Cao, X.Y., You, Y., Meng, Z.F., Sun, K.Y., and Shen, X.Y. (2023). Matrine suppresses NLRP3 inflammasome activation via regulating PTPN2/JNK/SREBP2 pathway in sepsis. Phytomedicine 109, 154574. 10.1016/j.phymed.2022.154574.

43. Wang, S., Zhou, Y., Yu, R., Ling, J., Li, B., Yang, C., Cheng, Z., Qian, R., Lin, Z., Yu, C. et al. (2023). Loss of hepatic FTCD promotes lipid accumulation and hepatocarcinogenesis by upregulating PPARgamma and SREBP2. JHEP Rep 5, 100843. 10.1016/j.jhepr.2023.100843.

44. Chen, W., Wen, L., Bao, Y., Tang, Z., Zhao, J., Zhang, X., Wei, T., Zhang, J., Ma, T., Zhang, Q. et al. (2022). Gut flora disequilibrium promotes the initiation of liver cancer by modulating tryptophan metabolism and up-regulating SREBP2. Proc Natl Acad Sci U S A 119, e2203894119. 10.1073/pnas.2203894119.

45. Foster, C., Gagnon, C.A., and Ashraf, A.P. (2024). Altered lipid metabolism and the development of metabolic-associated fatty liver disease. Curr Opin Lipidol 35, 200–207. 10.1097/MOL.0000000000000933.

46. Devang, N., Adhikari, P., Nandini, M., Satyamoorthy, K., and Rai, P.S. (2021). Effect of licorice on patients with HSD11B1 gene polymorphisms-a pilot study. J Ayurveda Integr Med 12, 131–135. 10.1016/j.jaim.2020.06.006.

47. Leyla Galandarli1*, C.M., Asli Javadzade1, Afat Mammadova1, , and Amrahov1, N. (2022). MOLECULAR DOCKING OF GLYCYRRHIZIN WITH 11β-HSD1 PROTEIN RESEARCH IN: AGRICULTURAL & VETERINARY SCIENCES 6, 141–146.

48. Hughes, K.A., Webster, S.P., and Walker, B.R. (2008). 11-Beta-hydroxysteroid dehydrogenase type 1 (11beta-HSD1) inhibitors in type 2 diabetes mellitus and obesity. Expert Opin Investig Drugs 17, 481–496. 10.1517/13543784.17.4.481.

49. Honma, T., Shinohara, N., Ito, J., Kijima, R., Sugawara, S., Arai, T., Tsuduki, T., and Ikeda, I. (2012). High-fat diet intake accelerates aging, increases expression of Hsd11b1, and promotes lipid accumulation in liver of SAMP10 mouse. Biogerontology 13, 93–103. 10.1007/s10522-011-9363-2.

50. Blaschke, M., Koepp, R., Streit, F., Beismann, J., Manthey, G., Seitz, M.T., Kragl, A., and Siggelkow, H. (2021). The rise in expression and activity of 11beta-HSD1 in human mesenchymal progenitor cells induces adipogenesis through increased local cortisol synthesis. J Steroid Biochem Mol Biol 210, 105850. 10.1016/j.jsbmb.2021.105850.

51. Morton, N.M., Holmes, M.C., Fievet, C., Staels, B., Tailleux, A., Mullins, J.J., and Seckl, J.R. (2001). Improved lipid and lipoprotein profile, hepatic insulin sensitivity, and glucose tolerance in 11beta-hydroxysteroid dehydrogenase type 1 null mice. J Biol Chem 276, 41293–41300. 10.1074/jbc.M103676200.

52. Lee, S.Y., Kim, S., Choi, I., Song, Y., Kim, N., Ryu, H.C., Lim, J.W., Kang, H.J., Kim, J., and Seo, H.R. (2022). Inhibition of 11beta-hydroxysteroid dehydrogenase 1 relieves fibrosis through depolarizing of hepatic stellate cell in NASH. Cell Death Dis 13, 1011. 10.1038/s41419-022-05452-x.

53. Maier, C.R., Hartmann, O., Prieto-Garcia, C., Al-Shami, K.M., Schlicker, L., Vogel, F.C.E., Haid, S., Klann, K., Buck, V., Münch, C. et al. (2023). USP28 controls SREBP2 and the mevalonate pathway to drive tumour growth in squamous cancer. Cell Death Differ 30, 1710–1725. 10.1038/s41418-023-01173-6.

54. Guo, C., Chi, Z., Jiang, D., Xu, T., Yu, W., Wang, Z., Chen, S., Zhang, L., Liu, Q., Guo, X. et al. (2018). Cholesterol Homeostatic Regulator SCAP-SREBP2 Integrates NLRP3 Inflammasome Activation and Cholesterol Biosynthetic Signaling in Macrophages. Immunity 49, 842–856.e847. 10.1016/j.immuni.2018.08.021.

55. Hu, M., Han, T., Pan, Q., Ni, D., Gao, F., Wang, L., Ren, H., Zhang, X., Jiao, H., Wang, Y. et al. (2022). The GR-gp78 Pathway is involved in Hepatic Lipid Accumulation Induced by Overexpression of 11β-HSD1. Int J Biol Sci 18, 3107–3121. 10.7150/ijbs.42376.

56. Zhang, Y., Zhu, Z., Sun, L., Yin, W., Liang, Y., Chen, H., Bi, Y., Zhai, W., Yin, Y., and Zhang, W. (2023). Hepatic G Protein-Coupled Receptor 180 Deficiency Ameliorates High Fat Diet-Induced Lipid Accumulation via the Gi-PKA-SREBP Pathway. Nutrients 15. 10.3390/nu15081838.

57. Xu, D., Wang, Z., Xia, Y., Shao, F., Xia, W., Wei, Y., Li, X., Qian, X., Lee, J.H., Du, L. et al. (2020). The gluconeogenic enzyme PCK1 phosphorylates INSIG1/2 for lipogenesis. Nature 580, 530–535. 10.1038/s41586-020-2183-2.

